# *In vitro* endoderm emergence and self-organisation in the absence of extraembryonic tissues and embryonic architecture

**DOI:** 10.1101/2020.06.07.138883

**Authors:** Stefano Vianello, Matthias P. Lutolf

## Abstract

The endoderm is the cell lineage which gives rise in the embryo to the organs of the respiratory and gastrointestinal system. Uniquely, endodermal tissue does not just derive from descendants of the embryo proper (the epiblast) but instead arises from their gradual incorporation into an extraembryonic substrate (the visceral endoderm). Given the configuration of the early embryo, such a paradigm requires epiblast endodermal progenitors to negotiate embryonic compartments with very diverse epithelial character, a developmental contingency reflected by the fact that key early endodermal markers such as *Foxa2* and *Sox17* have been consistently found to be embedded within gene programmes involved in epithelialisation.

To explore the underlying cell biology of embryonic endoderm precursors, and to explore the relationship between endoderm development, epithelial identity, and extraembryonic mixing, we leveraged Gastruloids, *in vitro* models of early development. These self-organising three-dimensional aggregates of mouse embryonic stem cells do not possess an extraembryonic component, nor do they appear to display typical tissue architecture. Yet, they generate cells expressing endodermal markers. By tracking these cells throughout *in vitro* development, we highlight a persistent and uninterrupted pairing between epithelial and endodermal identity, with FoxA2+/Sox17+ endoderm progenitors never transitioning through mesenchymal intermediates and never leaving the epithelial compartment in which they arise. We also document the dramatic morphogenesis of these progenitors into a macroscopic epithelial primordium extending along the entire anterior-posterior axis of the Gastruloid. Finally, we find that this primordium correctly patterns into broad domains of gene expression, and matures cells with anterior foregut, midgut, and hindgut identities within 7 days of culture. We thus postulate that Gastruloids may serve as a potential source of endodermal types difficult to obtain through classical 2D differentiation protocols.

## Introduction

In mouse and humans, the digestive track, the respiratory system, and key internal organs such as the thymus, the bladder, the pancreas and the liver all derive from the same progenitor tissue (Carlson, 2014; Lewis & Tam, 2006; Nowotschin et al., 2019a), a “mucous layer” first described in chick embryos (Pander, 1817), and which we now know as “endoderm” (Allman, 1854; Oppenheimer, 1940).

In mouse, this “inner skin” actually first assembles on the outer surface of the embryo, through a unique choreography of cellular movements illustrated in Figure 1 (Burtscher & Lickert, 2009; Kwon et al., 2008; Probst et al., 2021; Viotti et al., 2014). As the mouse embryo implants into the uterus of the mother, and extraembryonic tissues proliferate to impart to the conceptus its characteristic cylindrical shape (Smith, 1985), the mouse embryo (at the very tip of such cylinder) is little more than an epithelial mass of potent cells: the epiblast. Remarkably, such inconspicuous tissue will act as the origin of almost all cells of the developing embryo through the transformations brought about by gastrulation, key milestone of all embryonic development. As such, the initially multipotent and uncommitted cells of the early epiblast commit to specific fates, which are generally classified into the broad germ layer categories of ectoderm (skin and neural types), mesoderm (heart, muscles, and mesenchyme), and endoderm (internal organs, respiratory and digestive tract) (Arnold & Robertson, 2009; Takaoka & Hamada, 2012; Tam & Behringer, 1997; Tam & Loebel, 2007).

**Fig. 1.**
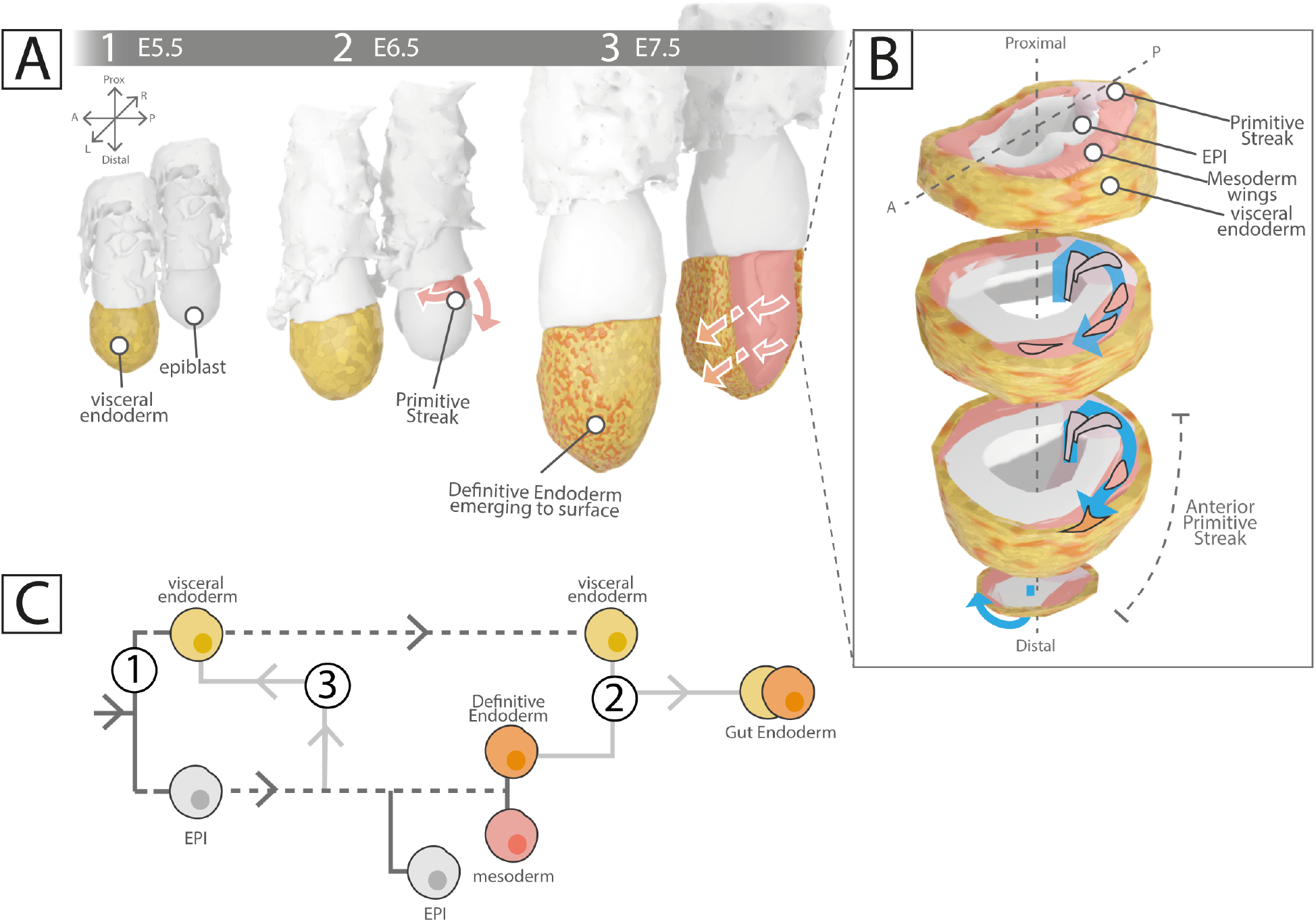
Endoderm development in the mouse embryo. (**A**) Intercalation of (embryonic) definitive endoderm cells (orange) into the visceral endoderm epithelium (yellow, extraembryonic), during peri-gastrulation stages of mouse development. Embryos in the back row are represented with their visceral endoderm layer removed. In red, mesodermal cells emerging from the primitive streak and starting their posterior-to-anterior circumnavigation of the epiblast. (**B**) Exploded view of the mouse embryo at around E7.5, as if sectioned proximo-distally. At the posterior of the epiblast a zone of EMT (primitive streak, light purple) advancing towards the disal tip of the embryo mediates epiblast cell egression. Cells specified to mesoderm (red cells) leave the epiblast and form so-called mesodermal wings as they circumnavigate the epiblast. They are sandwiched between the epiblast they just left (gray), and the overlaying visceral endoderm (yellow). Defintive endoderm cells (orange) transit only shortly within the mesodermal compartment, and instead egress into the visceral endoderm. (**C**) The multiple origins of endodermal cells, as adapted from (Nowotschin et al., 2019b). 1) visceral endoderm cells, originating from earlier segregation between “embryonic” and “extraembryonic” cell types in the blastocyst, 2) intermixing with definitive endoderm cells arising from gastrulation (see previous panels), 3) direct delamination from the epiblast prior to gastrulation. E = embryonic day.

Yet we now know that at least one of these germ layers, the endoderm, does not come entirely from cells originating in the epiblast (Kwon et al., 2008; Nowotschin et al., 2019a,b; Viotti et al., 2014). Blurring the boundaries of prevalent developmental paradigms, and making such qualifiers somehow oxymoronic, “extraembryonic” cells (i.e. cells that segregated away from the epiblast early on even before implantation) also contribute to the embryo, and specifically to its endoderm-derived tissues. Indeed, the epiblast is not isolated from other tissues within the conceptus, and it is actually enveloped by a thin epithelium of so-called Visceral Endoderm (Figure 1A, in yellow). This latter sheet of cells on the surface of the epiblast is what one would classify as extraembryonic tissue, originating from much earlier lineage segregation events (Chazaud et al., 2006). Originally thought to be displaced away as the embryo develops, this thin sheet of “extraembryonic” cells is the primary destination of a subset of cells produced by the epiblast (i.e. embryonic cells, in orange throughout Figure 1A), that these embryonic cells are those that will make endoderm, and as such that the visceral endoderm “extraembryonic” sheet in which these cells intercalate becomes itself an integrated component of later embryonic structures (Burtscher & Lickert, 2009; Kwon et al., 2008; Nowotschin et al., 2019b; Viotti et al., 2014).

What is the current model for how this process starts and unfolds? At around embryonic day (E)6.25 asymmetric signalling by extraembryonic tissues surrounding the epiblast break its symmetry (Stower & Srinivas, 2018; Takaoka & Hamada, 2012). One side of the epiblast starts becoming different from the rest (Figure 1A.2). At this side, which is defined as the posterior of the embryo and at which extraembryonic tissues concentrated high Wnt, BMP, and FGF signalling activity, epiblast cells respond by engaging so-called Epithelial to Mesenchymal Transition (EMT) programmes: they start losing attachment with the rest of the epithelium, they become motile and mesenchymal, they leave the epiblast (Arnold & Robertson, 2009; Tam & Behringer, 1997; Tam & Loebel, 2007). Morphologically, the so-called Primitive Streak appears: a distally-expanding zone of EMT leading to delamination of epiblast cells and simultaneous commitment to embryonic fates (Hashimoto & Nakatsuji, 1989; Williams et al., 2011). As epiblast cells undergo EMT and leave the epiblast, they start circumnavigating its outer surface, sandwiched under the overlaying visceral endoderm, forming wings of tissue converging towards the anterior of the embryo ((Hashimoto & Nakatsuji, 1989; Saykali et al., 2019; Viotti et al., 2014); red intervening intervening tissue in Figure 1A.2 and 1A.3, and red cell trajectory in Figure 1B). These cells will generate mesodermal derivatives, heart and muscles (Tam & Behringer, 1997).

Within the mesodermal compartment of the embryo another cell type finds its way: endodermal cells. Like mesodermal cells, these cells were once epiblast cells that left that epithelium to egress into the mesodermal compartment (Burtscher & Lickert, 2009; Probst et al., 2021). Rather than remaining within these wings of mesoderm however, endodermal cells start establishing contacts with the over-laying epithelium, the visceral endoderm, into which they eventually integrate (orange cell in Figure 11B; (Burtscher & Lickert, 2009; Kwon et al., 2008; Viotti et al., 2014)). The outer surface of the embryo thus quickly becomes a mosaic of its original resident population, that of visceral endoderm cells, and of an increasing number of ingressing and intercalating endoderm cells of embryonic (epiblast) origin, so-called Definitive Endoderm cells ((Burtscher & Lickert, 2009; Kwon et al., 2008; Viotti et al., 2014); orange cells in Figure 1A.3). This sheet of cells will later form pockets at the anterior and posterior of the embryo, and finally fold along its midline to close into a tube that will end up internalised within the embryo (not illustrated; (Carlson, 2014; Lewis & Tam, 2006; McGrath & Wells, 2015)). The gut tube has formed, and along its entire length progenitors of all endoderm-derived visceral organs will emerge and take shape (Carlson, 2014; McGrath & Wells, 2015).

Clearly then, what ultimately becomes the tissue we refer to as “gut endoderm”, i.e. the endodermal sheet which folds and closes to give rise to the embryonic gut tube, is thus actually a mixture of cells of very different origins, even though these converge towards similar (yet not identical) endpoint molecular signatures (Nowotschin et al., 2019a,b; Viotti et al., 2014). The multiple contributions to gut endoderm described above are summarised in Figure 1C (as adapted from e.g. (Nowotschin et al., 2019a)). In addition to the first contribution of Visceral Endoderm cells by early segregation within the inner cell mass of the early embryo (Figure 1C.1), and to the later intercalation of Defintive Endodem cells (Figure 1C.2), we also highlight a third source of cells: epiblast cells bypassing EMT and altogether bypassing transit within the mesodermal compartment of the embryo. These cells leave the epiblast to directly intercalate into the visceral endoderm, a contribution that has been documented to occur at the distal tip of the pre-gastrulation mouse embryo, zone of maximal mechanical stress (Hiramatsu et al., 2013; Matsuo & Hiramatsu, 2017), and that has found support from single-cell transcriptome analyses (Nowotschin et al., 2019b). Direct epiblast to endoderm transitions are particularly interesting, as even endodermal progenitors that do classically egress from the epiblast into the mesodermal space might do so by EMT processes different than those governing the egression and specification of mesoderm (Bardot & Hadjantonakis, 2020; Burtscher & Lickert, 2009; Probst et al., 2021), and the issue remains contentious. Currently, data suggests that egressing endodermal progenitors do not completely lose their epithelial character but instead transiently redistribute their surface adhesion molecules as they travel along the mesodermal compartment, until they contact their new epithelial niche, the visceral endoderm, and fully repolarise (Bardot & Hadjantonakis, 2020; Kwon et al., 2008; Nowotschin et al., 2019a; Viotti et al., 2014). Indeed, recent transcriptional comparisons have confirmed that endodermal progenitors show reduced expression of EMT and migration determinants compared to their mesodermal counterparts, suggesting separate and distinct modes of delamination (Probst et al., 2021)

Uncertainty remains regarding many of the steps described above, and on the exact nature of the transition states that embryonic endoderm precursors traverse as they leave epiblast potency and refine endodermal identity (Bardot & Hadjantonakis, 2020; Ferrer-Vaquer et al., 2010; Lewis & Tam, 2006). Notably, evidence for so-called “mesendodermal progenitors”, whereby bipotential cells able to give rise to both mesoderm and endoderm would exist within or outside the epiblast, is debated in mouse (Lewis & Tam, 2006) despite the clear existence of such progenitor state in other developmental models (e.g. sea urchin and roundworms; (Peter & Davidson, 2010; Sulston et al., 1983)). Certainly, segregation between the two germ layers in the mouse is documented already at very early stages, and actually within the very pre-streak epiblast (Burtscher & Lickert, 2009; Probst et al., 2021) and most recent explorations of the topic appear to indicate that endomesodermal progenitors do not stably arise during early endoderm development *in vivo* (Mittnenzweig et al., 2021; Probst et al., 2021). Uncertainty also remains on whether endodermal cells egress from the epiblast through mechanisms common to those of egressing mesodermal cells or through alternative mechanisms. Crucially, the transcriptional similarity between endoderm (but not mesoderm) progenitors with epiblast cells (Probst et al., 2021), the observation that endodermal cells can be seen to have left the epiblast in regions which the primitive streak has not yet reached (Burtscher & Lickert, 2009), and that those within mesoderm wings of the embryo have not lost their epithelial identity (Viotti et al., 2014) raise interest in the relationship between epitheliality and endodermal identity (Ferrer-Vaquer et al., 2010; Nowotschin et al., 2019a; Viotti et al., 2014). Given the recent spotlight on the mixed composition and distribution of gut endoderm cells (Nowotschin et al., 2019b), one also wonders whether extraembryonic and embryonic endoderm cells play distinct essential roles within this primordium and its derivatives. In embryos where embryonic endoderm precursors cannot integrate their extraembryonic substrate and remain trapped within the mesodermal compartment, these seem to lose their identity and embryos do not form midgut and hindgut (Kanai-Azuma et al., 2002; Viotti et al., 2014).

As a platform to explore the underlying cell biology of embryonic endoderm precursors, and to explore the relationship between endoderm development, epithelial identity, and extraembryonic mixing, we use Gastruloids (Beccari et al., 2018; Turner et al., 2017; van den Brink et al., 2014). These stem cell aggregates develop *in vitro* in times and patterns that are surprisingly but consistently reminiscent of *in vivo* embryonic development. While mainly characterised in terms of mesodermal and neuromesodermal development (van den Brink et al., 2020), they have been crucially also found to specify endodermal identities ((Anlaş et al., 2021; Beccari et al., 2018; Cermola et al., 2019; Pour et al., 2019; Turner et al., 2017; van den Brink et al., 2014; Veenvliet et al., 2020); see also Discussion). Work with this system reflects a paradigm whereby leaving cells to their own self-organisation exposes intrinsic cellular programmes and developmental modules that would be otherwise masked by the regulative context of normal embryonic development (Davies, 2017; Shahbazi & Zernicka-Goetz, 2018; Turner et al., 2016). In this perspective, the absence of typical embryonic architecture, compartmentalisation, and extraembryonic tissues, makes Gastruloids particularly suitable to study the relevance of these features to endoderm development.

We here highlight a persistent and uninterrupted pairing between epithelial and endodermal identity, with FoxA2+/Sox17+ endoderm progenitors never transitioning through mesenchymal intermediates and never leaving the epithelial compartment in which they arise. We also document the dramatic morphogenesis of these progenitors into a macroscopic epithelial primordium extending along the entire anterior-posterior axis of the Gastruloid, patterned into broad domains of gene expression. Finally, we show that Gastruloids appear to give rise to patterned mature endodermal identities corresponding to the entire spectrum of fates observed in the embryonic gut tube, with notable representation of anterior foregut, midgut, and hindgut types. Corollarily we also highlight a strong epithelial component in Gastruloids, and thus the spontaneous emergence *in vitro* of stratified architectures and germ layer compartmentalisation.

## Results

To investigate the emergence, dynamics, and patterning of endoderm progenitors *in vitro* we started by generating Gastruloids (Baillie-Johnson et al., 2015; Beccari et al., 2018; van den Brink et al., 2014). Accordingly, we aggregated 300 mouse embryonic stem cells of a TBra/Sox1 double reporter line (described in (Deluz et al., 2016)) whose output Gastruloids have been extensively characterised in published literature and for which we have documented expression of markers of all three germ layers (Beccari et al., 2018); see Materials & Methods). As expected, when 300 of these mouse embryonic stem cells are seeded in individual wells of a low-adhesion 96well plate and maintained in N2B27 medium, these sediment to the bottom of the well and aggregate together in the first 48h of culture to form a compact sphere with defined edges by 72h (Figure 2). A pulse of the glycogen synthase kinase (GSK) 3 inhibitor (***/***CHIR99021 (Chiron) is then applied as a trigger of “gastrulation” and as to mimic the increase in Wnt signalling experienced by cells of the posterior mouse epiblast. Accordingly, the aggregate breaks symmetry (Figure 2A, asterisk). Morphologically, the spherical 72h aggregate becomes oblong by 96h, and extends a long protrusion that grows over time (120h, 144h). This posterior protrusion is marked by *TBra* (Brachyury) expression, marker of the posterior primitive streak and of the embryonic tail bud, and found to similarly define the posterior of the Gastruloid (Beccari et al., 2018; Turner et al., 2017; van den Brink et al., 2014).

**Fig. 2.**
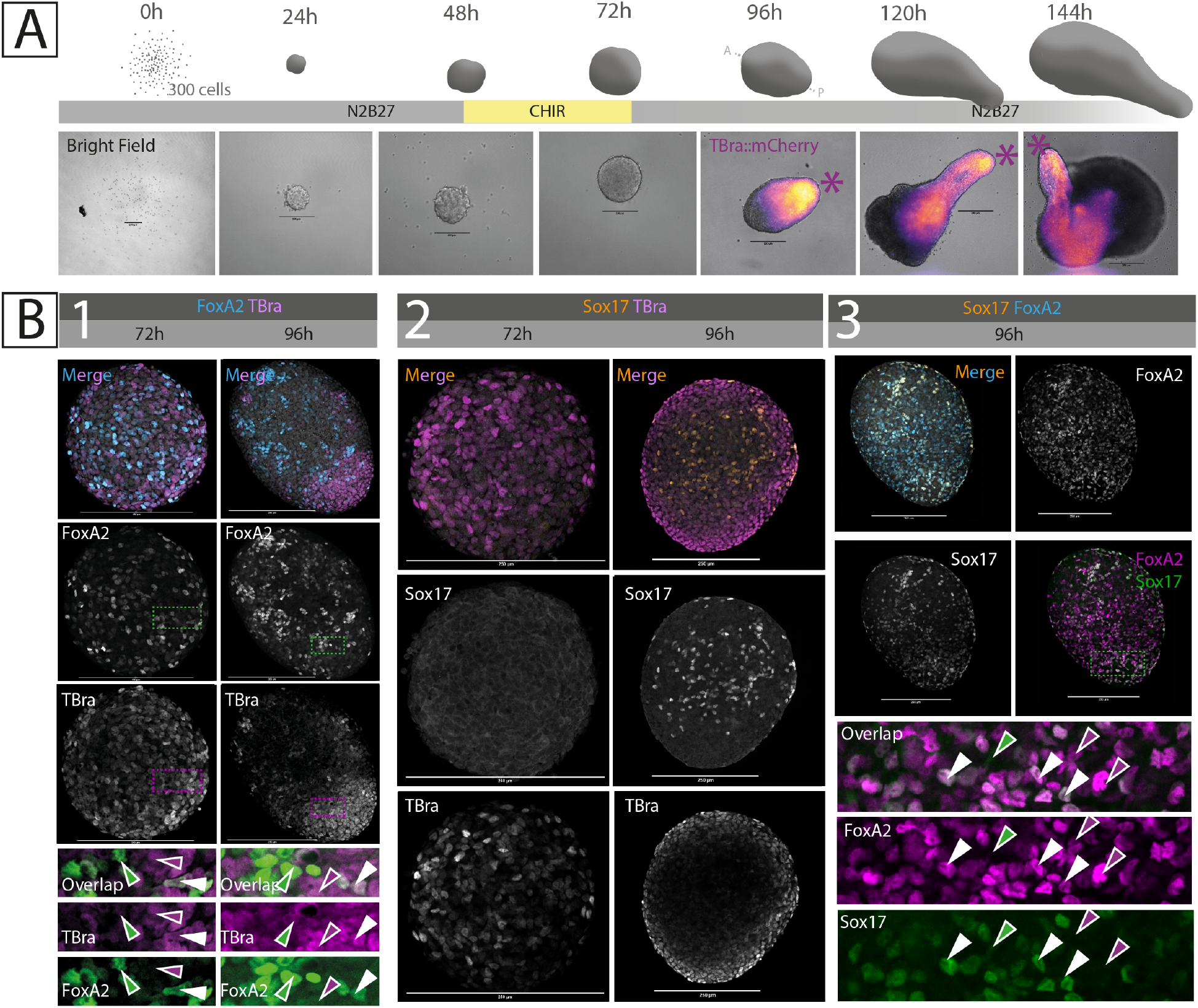
Emergence and patterning of endodermal markers in Gastruloids. (**A**) Summary schematic of the Gastruloid generation protocol (see Methods), and brightfield pictures of representative Gastruloids all along their development *in vitro*. Note that the fluorescence channel is here shown from t=96h onward only (when the Gastruloid becomes polarised). Reporter expression starts at t=72h, homogeneously throughout the spheroid (see (Turner et al., 2017)). (**B**) Immunostaining against the posterior epiblast and primitive streak marker TBra, and classical endodermal progenitor markers FoxA2 and Sox17, at 72h and 96h of Gastruloid development. SOX17+ cells appear one day later TBra+ and FOXA2+ cells, and are a nearly exclusive subset of the latter. Marker colocalisation is shown in green and magenta, with double-positive cells appearing white (examples of single-positive and double-positive cells highlighted by single-colour and white arrowheads respectively). Scale bar is always 250um. Asterisk indicates the posterior of the Gastruloid (based on TBra expression).

### Emergence and patterning of endodermal markers

#### Emergence of early endodermal markers

In the embryo, precursors of the definitive endoderm appear to be found within epiblast cells marked by expression of the transcription factor FOXA2 (Burtscher & Lickert, 2009; Probst et al., 2021). These FOXA2+ cells would be initially intermingled with TBra+ (TBXT) cells at the proximal posterior side of the embryo, and then resolve as a homogeneous FOXA2+ population marking the distal portion of the epiblast, and thus the epiblast region anterior to the leading edge of the primitive streak (Bardot et al., 2017; Burtscher & Lickert, 2009; Probst et al., 2021). While cells in the intervening epiblast region and co-expressing *FoxA2* and *TBra* may be progenitors of cardiac mesoderm types (Bardot et al., 2017), FOXA2+ cells of the distal epiblast are posited to leave the columnar epithelium, upregulate *FoxA2*, and move within the wings of mesoderm enveloping the epiblast ((Burtscher & Lickert, 2009; Kwon et al., 2008; Probst et al., 2021; Viotti et al., 2014), see also Figure 1B). FOXA2+ cells that contact the overlaying visceral endoderm would upregulate *Sox17* (Viotti et al., 2014), leave the mesodermal domain, integrate within this new epithelium, and join the cohort of cells that will eventually form the gut endoderm (Burtscher & Lickert, 2009; Kwon et al., 2008; Viotti et al., 2014). Given the relevance of *FoxA2* and *Sox17* for endoderm development (Dufort et al., 1998; Kanai-Azuma et al., 2002; Monaghan et al., 1993; Sasaki & Hogan, 1993), and their prevalent use in the gastruloid/embryoid literature as early endoderm markers (Beccari et al., 2018; Pour et al., 2019; Turner et al., 2017; van den Brink et al., 2014; Veenvliet et al., 2020), we decided to track their emergence and patterns of expression at the earliest stages of Gastruloid development.

At 72h, when the Gastruloid is still spherical and has just received the CHIR stimulus that will drive it through differentiation and morphogenesis, FOXA2+ cells can be seen scattered throughout the aggregate, intermingled with TBra+ cells (Figure 2B.1, 72h). Just 24h later (t=96h), and as TBra+ cells resolve into a pole that will accordingly define the posterior of the Gastruloid (van den Brink et al., 2014), FOXA2+ cells form clusters and segregate away from the TBra+ pole along the newly defined axis of the aggregate (Figure 2B.1, 96h). While few FOXA2+/TBra+ (double-positive) cells can be distinguished at this stage, most cells are either TBra+ at the posterior of the Gastruloid, or FOXA2+ as scattered clusters along the anterior (Figure 2B.1,bottom).

In contrast to TBra and FOXA2, SOX17 cannot be detected at t=72h, but only starts appearing later (Figure 2B.2). At t=96h, scattered unclustered SOX17+ cells appear within the elongating Gastruloid, and co-staining for FOXA2 shows these cells as representing a nearly exclusive subset of the FOXA2+ population (Figure 2B.3). We thus observe, at the earliest timepoints of Gastruloid response to CHIR, ordered emergence of key endodermal markers in sequence and patterns that are consistent with what is observed in the embryo. Not only SOX17+ cells emerge later and within a population of FOXA2+ cells (as seen in the embryo, (Viotti et al., 2014)), these cells sort from an initially TBra-intermingled population to later define posterior and anterior domains along the AP axis of the Gastruloid, just as is observed in the epiblast of the early primitive streak embryo (Burtscher & Lickert, 2009; Probst et al., 2021).

#### Cellular biology of endodermal cells

In vivo, FoxA2+ (and thus Sox17+) cells are expected to occupy and traverse very different embryonic compartments throughout their journey. As such, FOXA2+ cells would first emerge within the columnar epithelial tissue of the epiblast, they would then egress and mix with the mesenchymal mesodermal cell types circumnavigating the embryo as mesodermal wings, and they would finally re-integrate the epithelium on the surface of the embryo ((Kwon et al., 2008; Viotti et al., 2014), as illustrated in Figure 1B). Sox17 expression appears to be even more intimately associated with transitions between compartments, and has been reported to be expressed once endodermal precursors contact and integrate within the surface epithelium (Viotti et al., 2014). We thus sought to resolve whether cells expressing endodermal markers in Gastruloids were equivalently moving across compartments, with particular attention to their epithelial identity. Indeed, we observe SOX17+ cells in the absence of surface epithelial layer on which these would eventually integrate *in vivo*.

Co-staining for the the epithelial marker CDH1 (E-cadherin, adherens junction) shows that both TBra+ and FOXA2+ cells specified within the t=72h Gastruloid are emerging within a cellular aggregate that is uniformly epithelial (or, at least, epithelioid since epithelial architecture is missing, Figure 3A), consistently with the epithelial context of the epiblast of the early gastrulation embryo in which TBRA+ and FOXA2+ cells have been described to first emerge (Burtscher & Lickert, 2009; Lee et al., 2007; Probst et al., 2021). Cells of the t=72h Gastruloid all show homogeneous membrane CDH1 localisation, as likely expected for an aggregate of embryonic stem cells transitioning towards EpiSC states (Hamidi et al., 2020; Turner et al., 2017).

**Fig. 3.**
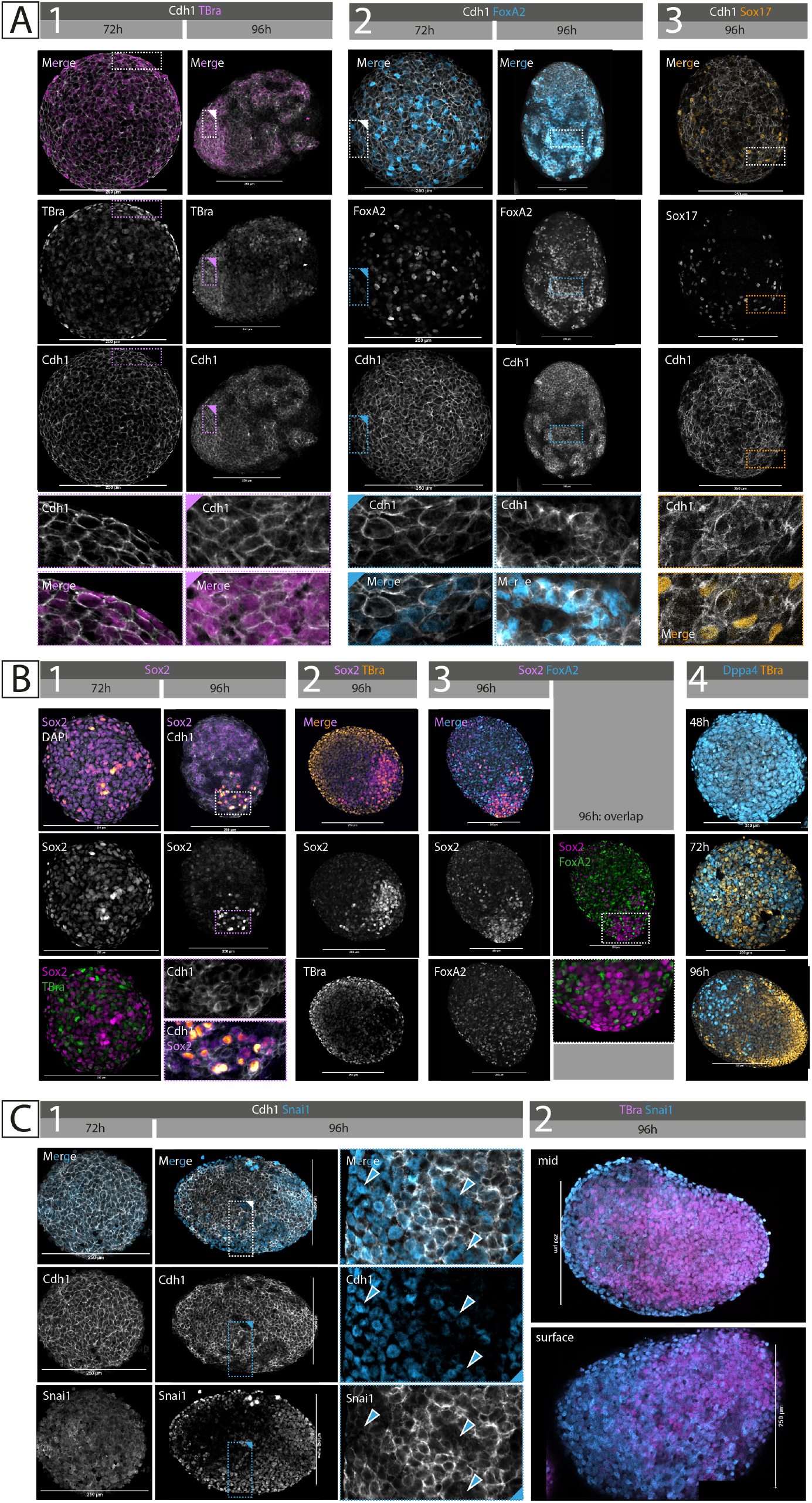
Endoderm progenitors are epithelial in nature, reside within an epiblast-like compartment, and are spared by classical EMT. (**A**) Immunostaining against the epithelial molecule E-cadherin (Cdh1) at 72h and 96h of Gastruloid development. Notice the drastic fragmentation of epithelial integrity between the two timepoints. Co-stained, are the posterior epiblast and primitive streak marker TBra, and classical endodermal progenitor markers FoxA2 and Sox17. Cells expressing either marker are consistently also CDH1+. (**B**) Immunostaining against the epiblast marker Sox2 shows its segregation at the anterior of the 96h Gastruloid, thus defining a separate domain from FoxA2+ cells. A similar pattern is highlighted by the pluripotency marker Dppa4. (**C**) Immunostaining against the classic EMT regulator Snail-1 (Snai1) shows patterns of expression complementary to those of CDH1. Sna1-mediated EMT is widespread throughout the 96h Gastruloid, and marks cells enveloping the aggregate. Scale bar is always 250um. Marker colocalisation is shown in green and magenta, with double-positive cells appearing white (examples of single-positive and double-positive cells highlighted by single-colour and white arrowheads respectively)

Interestingly, the CDH1 landscape of the t=96h Gastruloid, one day later, is radically different: as Gastruloids respond to the CHIR pulse, CDH1 expression becomes patchy. The original CDH1 continuum of the 72h spherical Gastruloid displays signs of clear fragmentation by t=96h (Figure 3A). At the posterior, CDH1+ cells remain clustered and maintain expression of TBra in a configuration analogous to that of the epiblast of the posterior or incipient primitive streak ((Burtscher & Lickert, 2009; Herrmann, 1991), Figure 3A.1), while anteriorly CDH1 continuity is increasingly interrupted by intervening non-epithelial cells (interpreted to be mesoderm). Very interestingly, all FOXA2+ cells seen at this stage are contained within this disaggregating CDH1 core, just as the newly specified SOX17+ cells emerging within such FOXA2+ population (Figure 3A.2 and Figure 3A.3). We never observe FOXA2+ or (FOXA2+/)SOX17+ cells outside of the perimeter defined by the CDH1+ islands.

These findings are particularly significant in that one might naively expect to observe FOXA2+ cells to be leaving their CDH1+ (“epiblast”) substrate and for FOXA2+ and FOXA2+/SOX17+ cells to be found in the emerging mesenchymal compartment. Significantly however, FOXA2+ cells leaving the epiblast *in vivo* do not lose CDH1 expression either, but rather relocalise it isotropically until they make contact with and reintegrate the overlaying visceral endoderm (Viotti et al., 2014). The isotropic CDH1 starting point of the Gastruloid at t=72h might thus reconcile *in vitro* and *in vivo* findings by explaining the retention of SOX17+ cells in the original “epiblast” compartment. Certainly, the observation of such pervasive CDH1 expression within Gastruloids, at least those derived from the stem cell line used in this study (see Discussion), challenged our pre-conception of Gastruloids as mainly mesenchymal organoids.

To test the underlying identity of the Cdh1 domain of t=72h and t=96h Gastruloids, and to check whether endodermal identities were indeed emerging and remaining associated with an “epiblast”-like domain, we co-stained Gastruloids for pluripotency and epiblast markers (Figure 3B). Staining for SOX2 shows that the t=72h spheroid is indeed a collection of SOX2+ cells (maintained from earlier timepoints, see data in (Beccari et al., 2018)), and newly emerging TBra+ cells (Figure 3B.1). As such the emerging scattered FOXA2+ population at this stage (also intermingled with TBra+ cells) is likely emerging on this very CDH1+/SOX2+ substrate. Yet, as gastruloid “gastrulation” progresses, the fragmenting epithelial compartment only maintains high SOX2 at a pole (opposite to the TBra pole, i.e. at the anterior, Figure 3B.2), such that the rest of the epithelium, where FOXA2+ and TBra+ identities segregate, shows low or no SOX2 (Figure 3B.3). A similar pattern of segregation and maintenance of potency at the anterior tip of the t=96h gastruloid is highlighted by the observed dynamics of the pluripotency marker DPPA4, marking the entire stem cell aggregate at t=48h, segregating from emerging TBra+ cells at t=72h, and being restricted at the anterior by t=96h (Figure 3B.4).

Our observations are consistent with a model where endodermal identities differentiate without ever leaving the epiblast compartment, or at least where endodermal precursors retain CDH1 expression throughout their development. Indeed, the CDH1+ mass of the t=96h Gastruloid may itself represent different embryonic compartments: one of epiblast, maintaining potency markers at the anterior and downregulating SOX2 at the posterior where FOXA2+ and TBra+ cells segregate to define respectively distal and proximal posterior epiblast identities, and one of FOXA2+/SOX17+ endodermal precursors that would normally be found in the mesenchymal compartment of the embryo but here remain attached to the “epiblast” given the isotropic CDH1 expression of all other compartments. In either case, we strengthen the case for direct epiblast-to-endoderm transitions that may not require classic EMT or transitions through so-called mesendodermal intermediates (Ferrer-Vaquer et al., 2010; Kubo et al., 2004; Lewis & Tam, 2006; Pfendler et al., 2005; Tada et al., 2005).

#### Gastruloids undergo widespread EMT, which spares endodermal precursors

To test whether the observed fragmentation of the epithelial core of the Gastruloid is consistent with EMT-like processes one would expect for an *in vitro* model of gastrulation, and how this relates to the apparent failed EMT of endodermal precursors in the Gastruloid, we performed immunostaining for the EMT master regulator Snail (SNAI1) ((Cano et al., 2000; Carver et al., 2001), Figure 3C).

While SNAI1 is only detected at low levels in the cytoplasm of cells of t=72h Gastruloids (all CDH1 positive, as shown before), large swathes of cells with strong nuclear SNAI1 signal are observed at t=96h (Figure 3C.1). Crucially, these patches of SNAI1+ cells are consistently observed to mark the cells intervening between fragments of the CDH1 core. Optical cross-sections at the midplane of t=96h Gastruloids show SNAI1+ cells forming an envelope at the surface, and establishing a posterior to anterior gradient in continuity with the TBra+ posterior (Figure 3C.2).

We thus notice that Gastruloid “EMT” seems to be a generalised rather than localised process, originating at several points within the CDH1+ “epiblast” substrate. Alternatively, SNAI1+ mesodermal types originating from a localised EMT origin may be migrating and physically displacing CDH1+ cells, leading to the observed fragmented appearance. Regardless, the retention of FOXA2+ cells within CDH1+ islands, in an environment of widespread EMT, seems to suggest that these cells are not leaving the epiblast and are either transitioning through endodermal differentiation within their original epiblast-like environment or attempting to leave the epithelium by Snai1-independent mechanisms (Bardot & Hadjantonakis, 2020; Probst et al., 2021) and either rapidly reintegrating it at short timescales or remaining attached to it through homotypic CDH1 interactions.

### Formation of an endoderm-like primordium

#### The epithelial core of the Gastruloids undergoes dramatic architectural rearrangements

To evaluate the later fate of endodermal cells within the gastruloid core, we further tracked Cdh1, FoxA2, and Sox17 patterns of expression as the Gastruloid undergoes morphogenesis and elongation from t=96h onward ((Beccari et al., 2018), Figure 4).

**Fig. 4.**
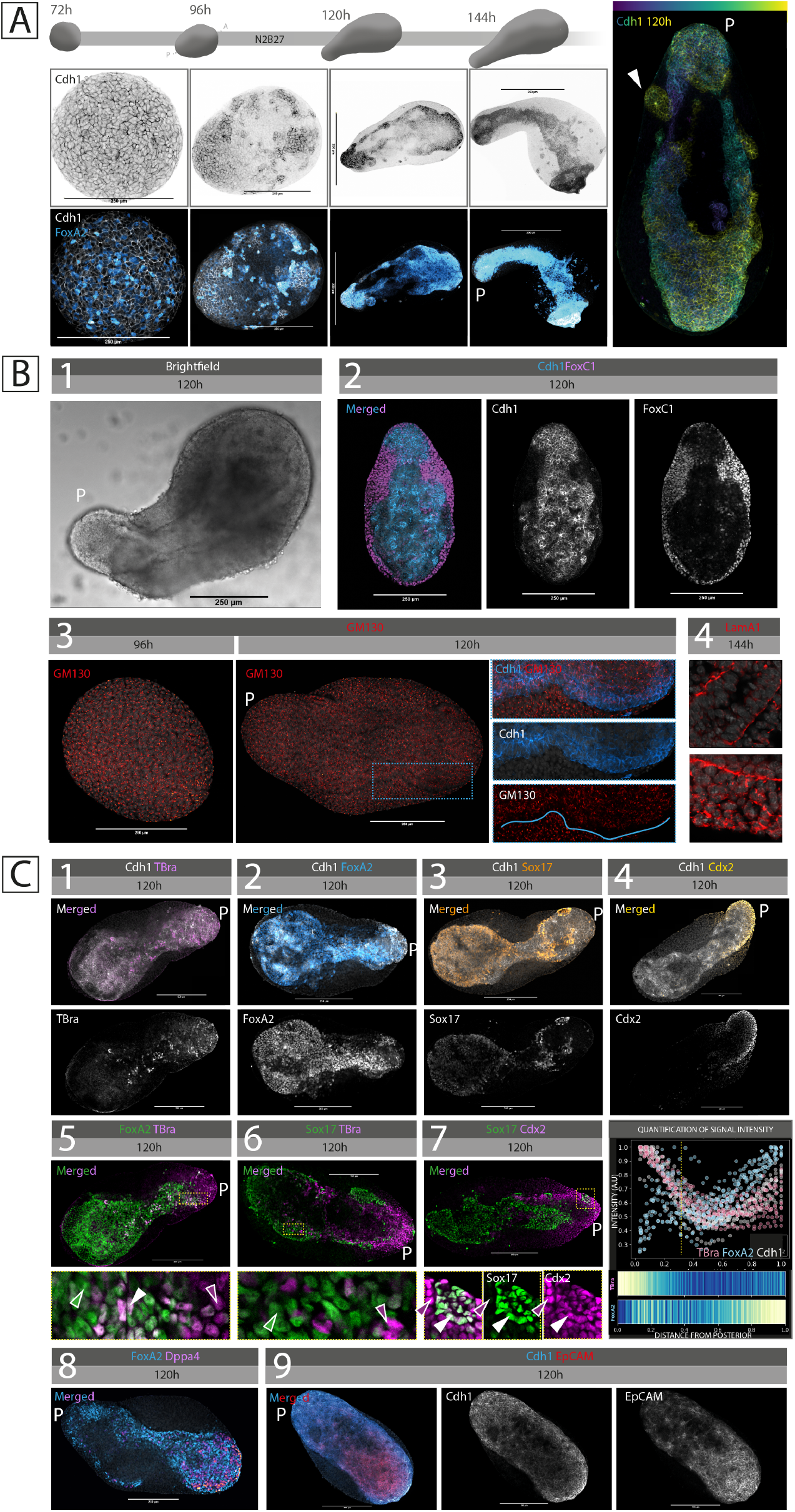
An epithelial primordium forms at the core of elongating Gastruloids, and shows patterned expression of endodermal markers. (**A**) Representative immunostainings of Gastruloids undergoing elongation and morphogenesis. CDH1+/FOXA2+ cells appear to segregate at the core of the Gastruloid and form an elongated primordium spanning the entire length of the aggregate. On the right, depth-coded projection of the primordium in a 120h Gastruloid. White arrowhead highlights a posterior lateral opening to the surface. (**B**) Macroscopic appearance of the epithelial primordium in its relationship with the overlaying mesodermal wings (FOXC1+). In the second row, maturation of the epithelial identity of the primordium is highlighted by the establishment of apico-basal polarity as hinted by GM130 segregation and basement membrane deposition (Laminin alpha 1 subunit, Lama1). (**C**) Immunostaining of elongated Gastruloids (120h) against the posterior markers TBra and Cdx2, the endodermal markers FoxA2 and Sox17, the pluripotency marker DPPA4, and the epithelial marker EpCAM to see their localisation within the CDH1+ primordium. A summary expression profile of some of these markers is also provided (gray dotted line indicating the separation between the domain of high TBra but low FoxA2, and that of high FoxA2 and low TBra).Scale bar is always 250um. Marker colocalisation is shown in green and magenta, with double-positive cells appearing white (examples of single-positive and double-positive cells highlighted by single-colour and white arrowheads respectively). P = posterior of the Gastruloid.

Strikingly, the fragmented CDH1+ core of the t=96h Gastruloid gradually re-organises in complex and streamlined elongated architectures extending along the entire length of the Gastruloid (Figure 4A). Over time, the tear-drop shaped, fragmented CDH1+ core of the t=96h Gastruloid tapers into a multi-branched, whisk-shaped epithelial primordium (t=120h, 144h), which in turn resolves in a single rod-like tissue that follows the outer geometry of the Gastruloid (144h onward). This epithelial structure, consistently seen in all (n = 97/99 imaged Gastruloids, N = 6 independent experiments) samples (Figure 5), emerges on the surface of the Gastruloid at the very posterior (and is indeed an extension of it), sometimes displaying multiple surface points in the region (white arrowhead Figure 4A, right), and occasionally also resurfaces at the anterior at much later timepoints (t=168h). Crucially, cells of this epithelial core are marked by FoxA2 at all stages of development (Figure 4A, bottom row). Macroscopically, this epithelial mass is already distinguishable in brightfield as a rod-like structure of compact cells extending from the posterior and gradually becoming enveloped by anteriorly-extending wings of loser, mesenchymal-like cells (Figure 4B.1). Co-staining of CDH1 with the pan-mesodermal marker FoxC1 (Sasaki & Hogan, 1993) clearly identifies the enveloping tissue as mesodermal (whose differentiation will give rise to the variety of trunk and cardiac structures described in (Rossi et al., 2019; van den Brink et al., 2020; Veenvliet et al., 2020)) and highlights a multilayered architectural organisation of the Gastruloid model, with interfacing epithelial and mesenchymal tissue (Figure 4B.2). Interestingly, such coupled configuration raises the possibility that the two compartments may engage in productive developmental interaction, possibly stimulating the development of cells types that would not otherwise emerge in either alone (as described *in vivo* e.g. in (Han et al., 2019)).

**Fig. 5.**
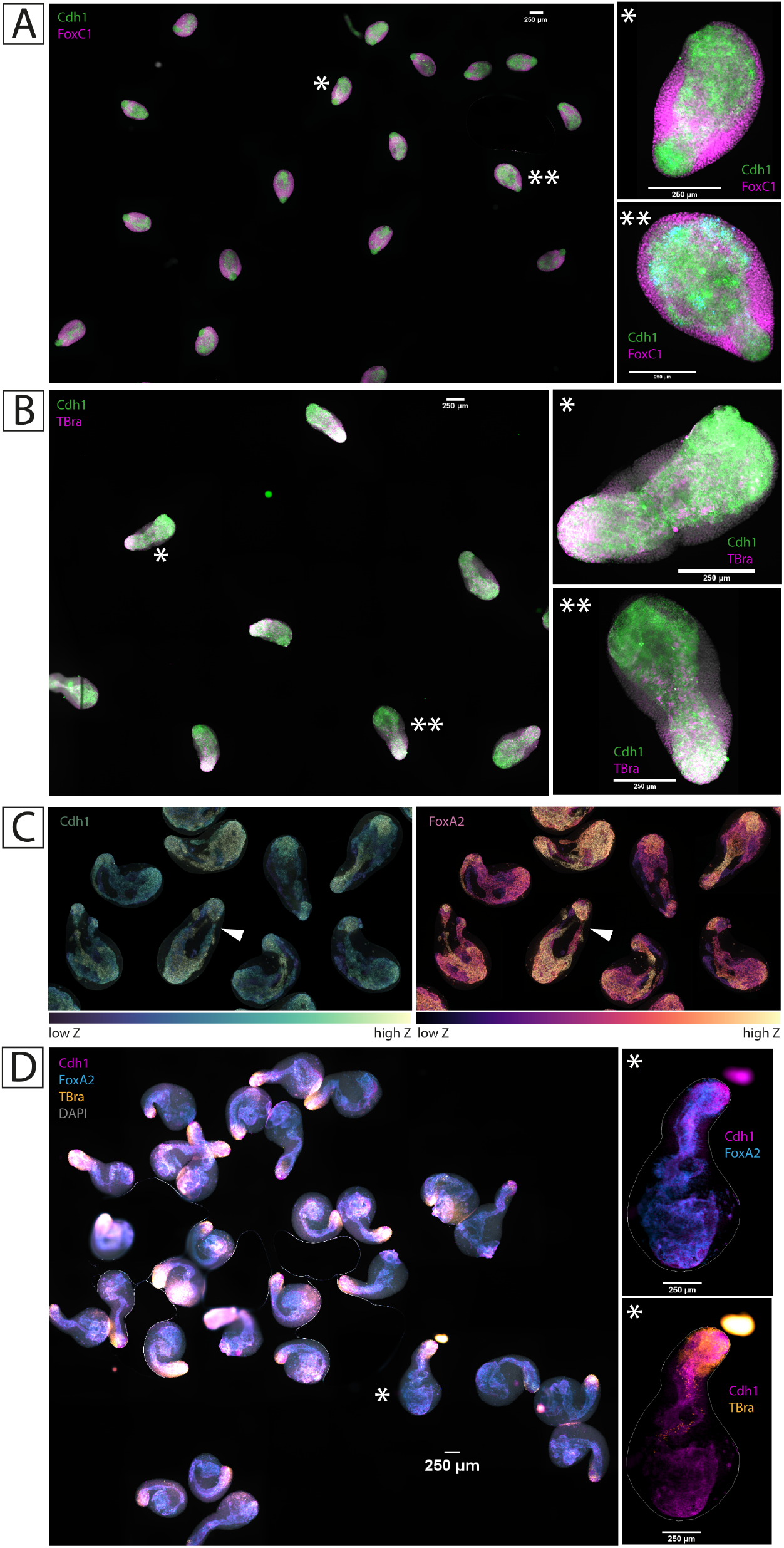
The CDH1 epithelial primordium is consistently observed in all TBra/Sox1 double reporter Gastruloids (n=97/99, N=6). Automated scanning of an entire microscope slide of t=120h Gastruloids, immunostained against CDH1 and either (**A**) the pan-mesodermal marker FOXC1, or (**B**) the posterior epiblast and primitive streak marker TBra. Asterisks indicate samples highlighted at the right of each panel. (**C**) Depth-coded collection of multiple t=120h Gastruloids, immunostained for CDH1 and FOXA2. White arrowhead indicates the sample shown in Figure 4A, right panel. Gastruloids from different regions of a same slide were here digitally placed close to each other. (**D**) Automated scanning of an entire microscope slide of t=144h Gastruloids, immunostained against CDH1, FOXA2, TBra. The asterisk indicates the sample highlighted at the right. Scale bar is always 250um.

To confirm the spatial dynamics of endoderm cells, we tracked live *FoxA2*+ cells by imaging Gastruloids formed by aggregation of FoxA2/TBra double reporter cells (TFoxA2; T/Bra:GFP, FoxA2:tagRFP, Yang (2015)). Gastruloids made from this cell line form too a CDH1+/FOXA2+ internal primordium by 144h, analogous to that observed in SBR Gastruloids (Figure 6A and Figure 6B). Live imaging of these Gastruloids shows *FoxA2*+ cells initially exclusively within the posterior (*TBra*+) domain of the Gastruloid. These cells then gradually migrate out of this domain to populate the anterior half of the Gastruloid where they proliferate and coalesce into the final compact primordium (Figure 6C, and Supplementary Video 1). The formation of a single, compact mass of cells by 144h thus appears the result of cells clustering to each other as they move from posterior to anterior, and as they move and divide locally within the anterior domain. FACS analysis of the double-reporter Gastruloids confirms the drastic increase in FoxA2+ cells from 96h and 120h (Figure 6D).

**Fig. 6.**
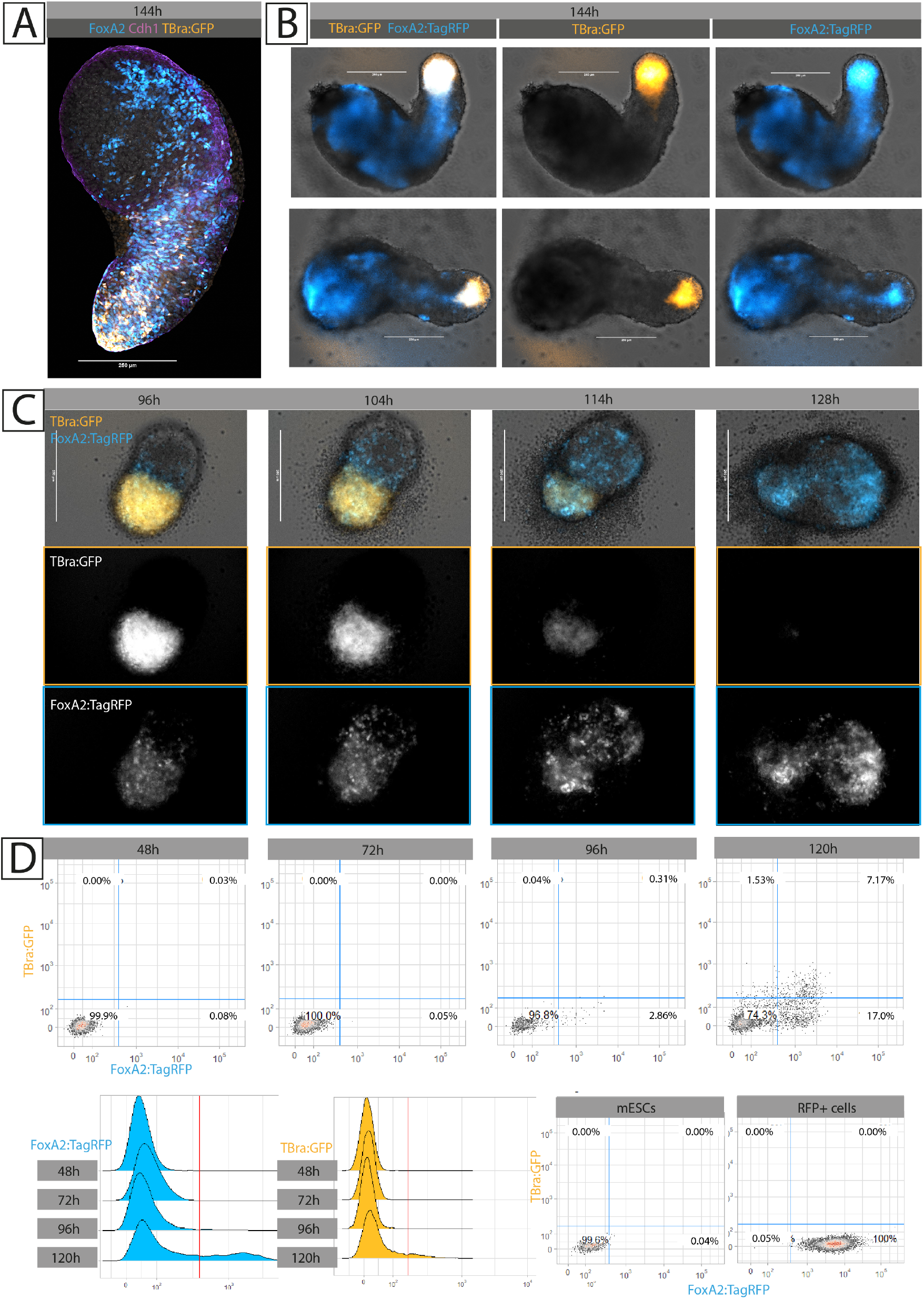
Gastruloids made from TBraGFP/FoxA2tagRFP cells highlight dynamics of endoderm primordium morphogenesis. (**A**) t=144h Gastruloid immunostained against FOXA2 (cyan) and CDH1 (magenta). The endogenous GFP signal (gold) shows cells positive for *TBra*. As in SBR Gastruloids, FOXA2+ cells populate a central epithelial domain. (**B**) Live imaging of reporter expression in t=144h Gastruloids shows a pattern of RFP (FoxA2, cyan) expression extending along the entire axis od the Gastruloid, consistent with what characterised in SBR Gastruloids (**C**) Still frames of a timelapse of Gastruloid development from t=96h to t=144h. RFP+ cells (FoxA2, cyan) emerge within the posterior domain of the Gastruloid (marked by TBraGFP expression, gold), populate the anterior domain, and coalesce and proliferate to form the internal rod-like primordium (**D**) FACS quantification of RFP+ cells over Gastruloid development shows a 8 fold increase between 96h and 120h. Thresholds were calibrated based on signal intensities in 2D-cultured double reporter stem cells (negative reference), and constitutively-expressing RFP+ cells (positive reference). Scale bar in all microscopy images is 250um.

Interestingly, the CDH1+ primordium appears to undergo epithelial maturation over time. Whereas CDH1+ cells of the t=72h Gastruloid represented more an epithelioid state, with expression of epithelial markers but without epithelial architecture, the t=120h and onward CDH1 mass shows signs of apico-basal polarity with polarised arrangement of GM130 (Figure 4B.3), and gradual deposition of discontinuous stretches of laminin (LAMA1 subunit) at the interface with the overlaying mesoderm (Figure 4B.4). We could not prove the existence of a continuous lumen within this epithelium, suggesting this structure to be more akin to a plastic epithelial mass rather than a defined continuous tube (data not shown). When present, supernumerary points of contact with the posterior surface of the Gastruloid do however seem to be consistently associated with rosetting and local invagination (see Figure 4A), even though we cannot at this point demonstrate continuity of these cavities with the rest of the CDH1 core.

#### Endodermal patterning along the AP axis of the Gastruloid

Having observed such dramatic and unexpected epithelial rearrangements over late Gastruloid development, and such tight association between this newly formed CDH1 core and the endodermal marker FOXA2 (Figure 4A and 5C), we proceeded to further characterise the identity of these cells and their possible patterning along the AP axis (Figure 4C, indeed observed for cells of the overlaying mesodermal compartment (Beccari et al., 2018; van den Brink et al., 2020)).

Matching the surprising increase in the number of CDH1+/FOXA2+ cells from t=96h to t=120h, where FOXA2 seems to mark almost the entirety of the CDH1+ primordium, SOX17 immunostaining also reveals a surprising increase in SOX17+ numbers, with cells extending from the neck of the CDH1 primordium (and often from the “hole” like surface openings described above) up to anteriormost extremity of the epithelium (Figure 4C.2 and Figure 4C.3). Immunostaining for the posterior primitive streak and tailbud marker TBra (Figure 4C.1) reveals that while FOXA2+ and SOX17+ cells may well be in continuity with the CDH1+/TBRA+ Gastruloid tip, they organise themselves just anterior to it. Batch quantification of signal intensity along the AP axis of Gastruloids (see plot in Figure 4C) indeed highlights reproducible patterning where the posterior 5th of the CDH1 primordium is marked by TBra+/CDH1+/FOXA2-cells, while the rest is populated by TBra-/CDH1+/FOXA2+ (and SOX17+) cells, an epithelialised endoderm whose continuity with the TBra+ pole might be explained by persistent homotypic interaction between different CDH1+ tissues. Of note, few TBra+ cells do seem to also extend deeper into the FOXA2+ domain (Figure 4C.5), and constitute a CDH1+/TBra+/FOXA2+ population that may be consistent with midline embryonic structures (Burtscher & Lickert, 2009; Yamanaka et al., 2007) (and captured by FACS, see Figure 6D).

At the very posterior of the Gastruloid (and thus of the CDH1 primordium), the posterior marker CDX2 (Beck et al., 1995) marks not only the TBra+/CDH1+ cells of the gastruloid tip, and CDH1-mesenchymal cells emerging laterally from it (Figure 4C.4), but also (CDH1+/)SOX17+ cells at the posterior limit of the SOX17+ domain (Figure 4C.7). These structures have been likened to the caudal intestinal portal forming during *in vivo* endoderm development (Beccari et al., 2018). At the opposite end (anterior), DPPA4+ cells intermingle with FOXA2+ endoderm (Figure 4C.8), possibly representing a surprising maintenance of pluripotency from the earliest timepoints of Gastruloid differentiation (giving their continuity with DPPA4+ cells at all previous timepoints). On this regard, other groups have interestingly reported the presence of Primordial-Germ-Cell-like cells, marked by DPPA3+, in association with the endodermal component (Veenvliet et al., 2020).

Finally, we identify the cell surface protein EpCAM as another marker of the entire primordium, with expression almost completely overlapping that of CDH1 (Figure 4C.9), yet with an apparent enrichment towards the anterior. The expression of EpCAM distinguishes the (CDH1+/)SOX17+ cells we observe as being indeed endodermal, given that this same marker also characterises endothelial progenitors (which would however be EpCAM-) at around the same developmental timepoints (Choi et al., 2012). Interestingly, EpCAM staining appears enriched in the region of the CDH1 primordium occupied by SOX17+ cells, hinting that combinations of cell-surface markers might drive further sub-sorting of different epithelial combinations within this same CDH1+ core. Here, the posterior of the primordium would represent a “posterior epiblast”-like, CDH1+/EpCAMlow/TBra+ domain; and the anterior a CDH1+/EpCAMhigh/FOXA2+/Sox17+ “endoderm”-like domain.

#### Gastruloid endoderm contains patterned anterior and posterior endodermal types

To better characterise the cell identities represented within the Gastruloid endoderm primordium, beyond immunostainings for classical markers, we made use of a recently released single-cell RNA sequencing dataset of SBR Gastruloids spanning timepoints t=96h to t=168h (Rossi et al., 2019). Analysis of the dataset highlights two clusters characterised by the expression of *FoxA2, Sox17, Cdh1*, and *EpCAM*, and lower numbers of cells also expressing *TBra* and *Eomes* (see Figure 7A, lower), and as such interpreted as to represent endoderm. Cells of one of such clusters are marked by the expression of genes such as *Otx2, Sox17, Hhex, Gata6, Gsc* (cluster 13 in Figure 7A, light yellow), and were here labelled as “early endoderm” given that these cells present a signature of anterior mesendoderm and definitive endoderm (Costello et al., 2015; Thomas et al., 1998). The second cluster was instead demarcated by the expression of genes such as *Cldn4,6,7, Krt7, Crb3* (cluster 4 in Figure 7A, beige) and was here labelled as “mature endoderm”, given the strong expression of epithelial markers shown to characterise later (rather than early) stages of endoderm maturation *in vivo* (i.e. gut tube; Anderson et al. (2008); Ogaki et al. (2011)). Indeed, performing differential expression analysis between the two clusters (Figure 7A, side-by-side grid) reveals that the “early endoderm” can be distinguished by higher expression of genes such as *Sfrp1, Lhx1, Hesx1, Fgf5* (consistent with an anterior endoderm/mesendoderm character; Costello et al. (2015); Finley et al. (2003); Khoa et al. (2016)), while “mature endoderm” distinctively expresses higher levels of *e*.*g. Igfbp5* and *Frem2* (expressed in the gut tube, Green et al. (1994); Timmer et al. (2005)), as well as additional epithelial markers such as *Krt19* and *Cldn4,9*. Adding a third dimension to the UMAP embedding representation (3D UMAP, Figure 7B) reveals that the two “endoderm” clusters reside in closest proximity to the cells annotated as anterior mesoderm (cluster 3; markers: *Gata6, Hand2, Myl7, Gata4, Lhfp*; in continuity with the “early endoderm” cluster 13), and ectoderm (cluster 5; markers: *Gjb3, Epcam, Tfap2c, Cdh1, Krt18*; in continuity with the “mature endoderm” cluster 4), but also with those annotated as notochord/axial mesoderm (cluster 1; markers: *TBra, Cobl, Shh, Cdx2, Noto*; also closest to the “mature endoderm” cluster 4). A list of all markers is available in as a Supplementary Table. Supporting the early/mature labelling of each cluster, the “early endoderm” cluster preferentially contains cells of Gastruloids from earlier timepoints (96h-144h), while the “mature endoderm” preferentially contains cells from later timepoints (120h-168h, Figure 7C). Notably, the “mature endoderm” cluster expresses all markers recovered by immunostaining (see previous figures and Figure 7A, lower panels), and many markers associated with foregut and anterior foregut (e.g. *Nkx2*.*3, Isl1, Otx2*; Biben et al. (2004); Nowotschin et al. (2019b); Zhuang et al. (2013)) (Figure 7D). Still, markers identifying all positions along the AP axis of the embryonic gut tube (Nowotschin et al., 2019b) could be recovered within Gastruloid endoderm clusters (Figure 7D and Figure S1).

**Fig. 7.**
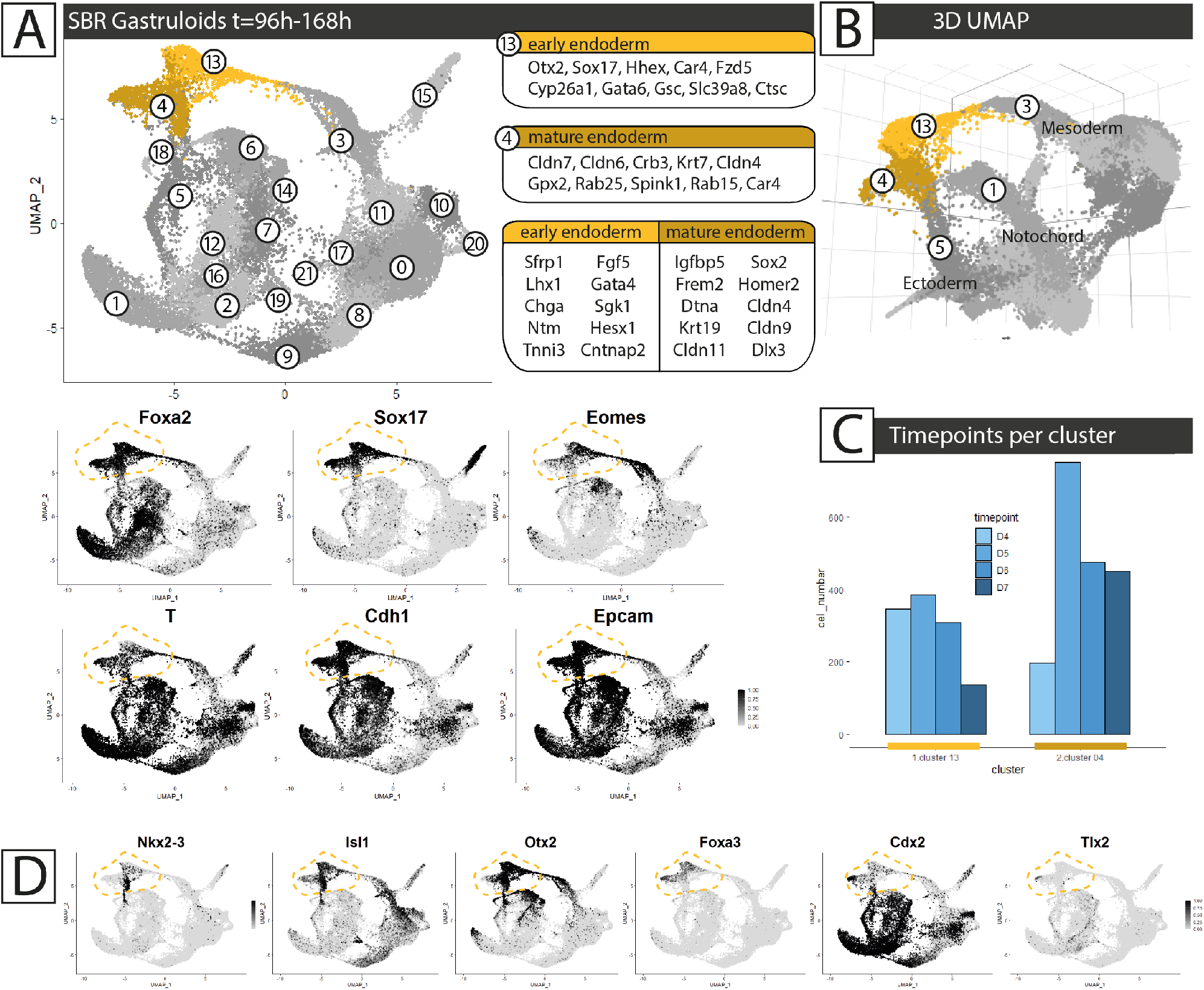
Analysis of single-cell RNAseq datasets from SBR Gastruloids highlights two “endoderm” clusters. (**A**) **Top:** UMAP representation of the dataset from Rossi et al. (2019). The two clusters attributed as “early endoderm” (13) and “mature endoderm” (4) are highlighted in gold and beige respectively. The top 10 marker genes for each cluster are indicated, as well as the top 10 differentialy expressed genes distinguishing one from the other. **Bottom**: Expression of classic endoderm markers (black). (**B**) 3D UMAP of the same dataset as in A. Clusters in proximity to the endoderm cluster are numbered, along with their annotation. Notice how cluster 1 appears distant in the 2D UMAP. (**C**) Distribution of cells from Gastruloids from each timepoint (Day4 to Day7, D4 to D7, i.e. 96h to 168h) across the two “endoderm” clusters. (**D**) Expression of anterior and posterior (left to right) gut tube markers (see Nowotschin et al. (2019b)) within the gastruloid dataset. “Endoderm” clusters are circled in gold throughout.

To better resolve the extent to which gut tube endoderm identities are represented amongst Gastruloid endoderm, we referred to published single-cell RNAseq data of the mouse embryonic gut tube (Nowotschin et al., 2019b). The questions arises on whether endoderm cells making up the central core of the Gastruloids represent the entirety of the embryonic gut tube or only a subset of it (e.g. only posterior endoderm), and whether the absent contribution of extraembryonic endoderm cells in the Gastruloid generation process may bias such identities to specific domains. More generally, we were interested in resolving the type of endoderm generated by self-organisation within Gastruloids. As in the original publication (Nowotschin et al., 2019b), the UMAP representation of embryonic endoderm cells (shown in Figure 8A and throughout) can act as an approximate visual map of the gut tube as it extends from anterior foregut (leftmost in Figure 8A) to posterior hindgut (up-right in Figure 8A). Accordingly, plotting known markers of anterior (e.g. *Nkx2*.*5, Sox2*; Nowotschin et al. (2019b); Wei & Condie (2011); Wood & Episkopou (1999); Zhang et al. (2005)) and posterior gut (e.g. *Cdx2, TBra*; Beck et al. (1995); Kispert & Herrmann (1994)) marks corresponding left and right regions in the UMAP (Figure 8A), and indeed plotting all 20 Transcription Factors identified by Nowotschin et al. (2019b) as marking increasingly posterior regions of the gut tube marks increasingly rightmost domains of the UMAP in our reprocessed dataset too (see Figure S1).

**Fig. 8.**
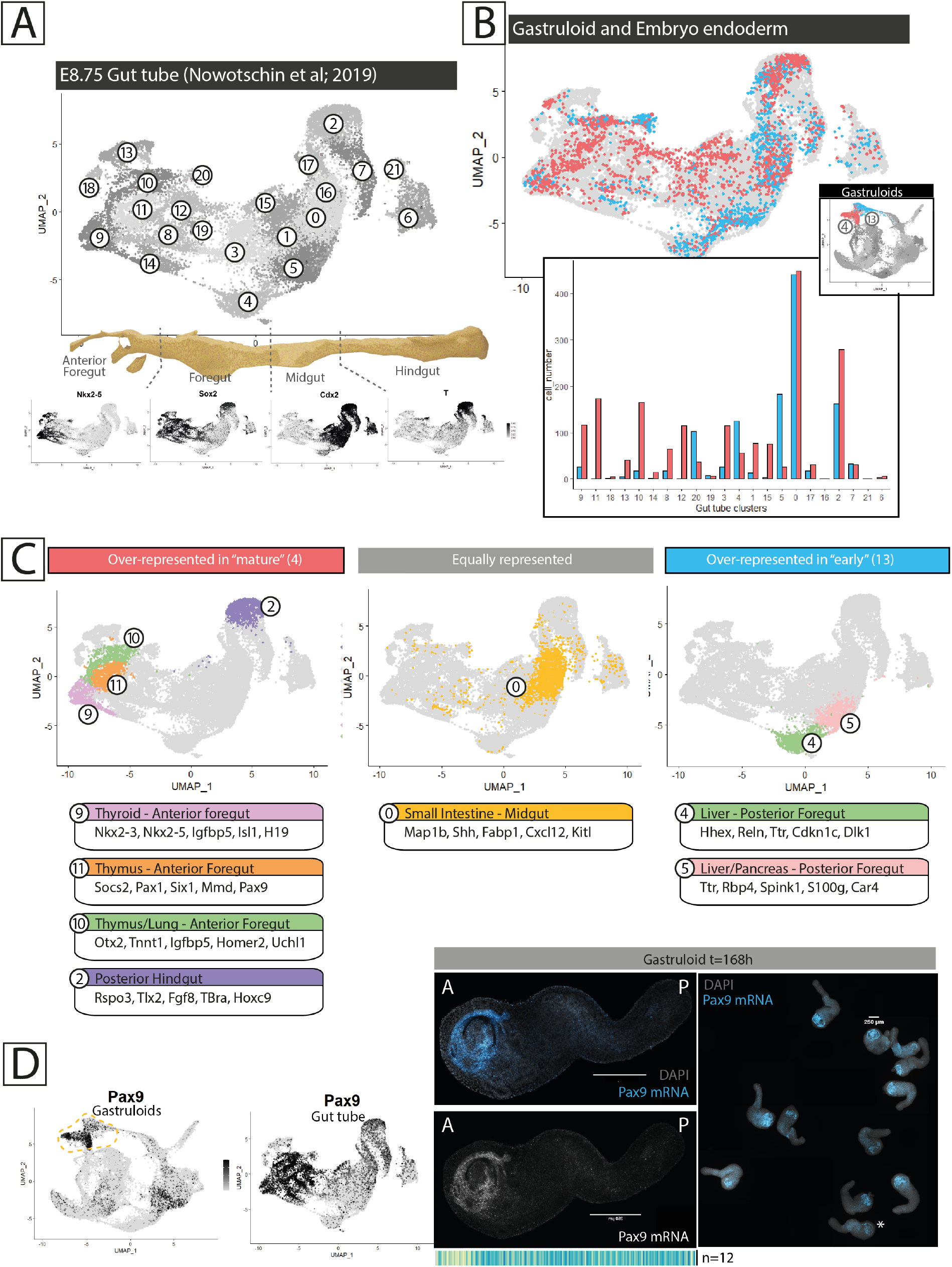
Gastruloid “mature endoderm” aligns with identities across the entire length of the embryonic gut tube, including Anterior Foregut. (**A**) UMAP of the reprocessed embryonic gut tube dataset from Nowotschin et al. (2019b). The expression domains of known anterior and posterior gut markers orients the map with anterior foregut at the left, and posterior hindgut at the right. (**B**) **Top:** Integration with batch correction of Gastruloid-endoderm clusters (4 and 13, red and blue respectively, see inset) and the embryonic gut tube reference (light gray). **Bottom:** barplot quantifying the number of cells from each gastruloid-endoderm cluster falling within each embryonic gut tube cluster (ordered from anteriormost to posteriormost left to right along the x-axis). (**C**) Highlight of the embryonic clusters populated majoritarily by Gastruloid-”mature endoderm” cells (**left**), “early endoderm” (**right**), or both (**middle**). The top 5 marker genes for each of these clusters is indicated, as well as their annotation. (**D**) Expression pattern of the Anterior Foregut marker *Pax9*, and corresponding Hybridisation Chain Reaction against *Pax9* mRNA in t=168h Gastruloids. Asterisk indicates the Gastruloid shown in the magnification. Scale bar is 250um throughout.

**Fig. 9.**
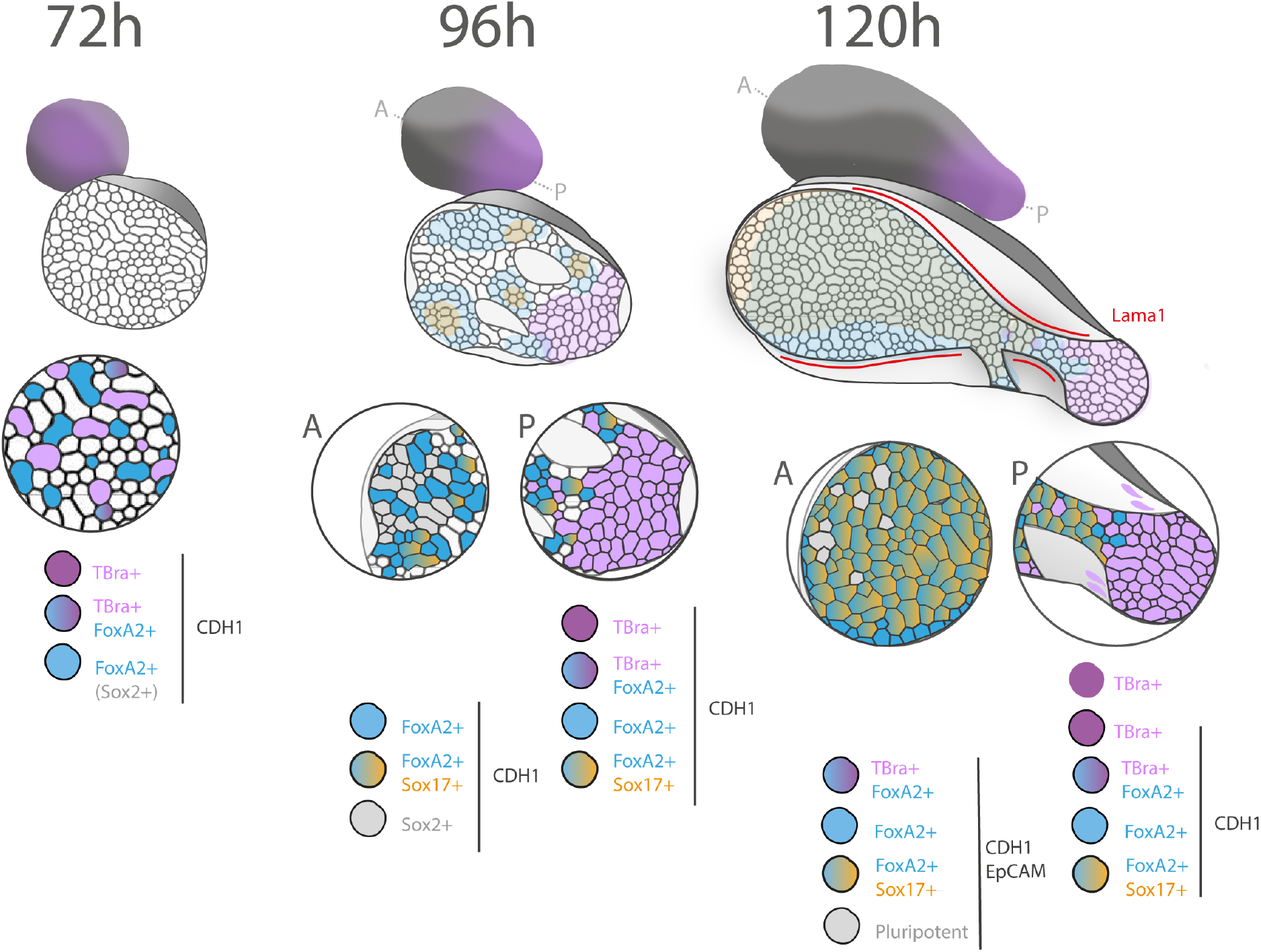
Endoderm specification and patterning in Gastruloids, a summary. At 72h of development, Gastruloids appear to be an epithelioid spheroid with intermingled TBra+ and FOXA2+ CDH1+ cells. We interpret FOXA2+ cells to also be SOX2+, and this spheroid to represent the posterior epiblast of the early gastrulating mouse embryo, with equally intermingled populations. Between 72h and 96h, Gastruloids undergo widespread Snai1-dependent EMT: mesenchymal types emerge and CDH1 continuity fragments. At the posterior, CDH1+/TBra+ cells define a proximal posterior epiblast compartment and FOXA2+ cells define a distal posterior epiblast region. Anteriorly, the CDH1 domain remains SOX2+. Within the CDH1+/FOXA2+ domains, SOX17+ cells emerge. In the embryo these cells would be expected to be found in the mesenchymal compartment. At t=120h, the CDH1+ primordium organises as a maturing epithelium extending along the entire length of the aggregate, enveloped by mesenchymal (mesoderm) cells. FOXA2+ and SOX17+ cells pattern along this primordium, just anteriorly to a posterior-epiblast-like TBra+/CDH1+ domain. EpCAM also seems to mark this very CDH1 primordium, with an anterior enrichment.

Leveraging the availability of this embryonic dataset, we integrated Gastruloid endoderm clusters (*in vitro* cells) and gut tube embryonic clusters (*in vivo* reference cells) with batch correction, to see how Gastruloid endoderm cells would distribute across the shared low dimensionality embedding (Tan et al., 2021). As shown and quantified in Figure 8B, cells from both Gastruloid endoderm clusters (“early”, cluster 13 in light blue, and “mature”, cluster 4 in red) span the entire length of the embryonic domain (light gray), with “early endoderm” cells showing a clear preferential accumulation within areas covered by cells of the posterior gut. The relative representation of “early” and “mature” Gastruloid endoderm cells within each region of the UMAP (Figure 8B, barplot) shows that i) “mature endoderm” cells from the Gastruloids span the entire length of the UMAP, and ii) they are the major Gastruloid-endoderm type falling within left-most clusters (Figure 8C, clusters 9, 10, 11). These clusters, within the gut tube dataset, would be annotated as anterior foregut types, progenitors of the thyroid, thymus, and lungs (Figure 8C, leftmost). Gastruloid-”mature endoderm” cells also fall within the embryonic Posterior Hindgut cluster (cluster 2) and, in equal proportions with “early endoderm” cells, within the cluster that would be annotated as Small Intestine (cluster 0, Figure 8C, middle). “Early endoderm” Gastruloid cells seem instead to over-represent Posterior Foregut/Midgut identities (Liver, Pancreas; clusters 4 and 5, Figure 8C, right). In summary, it appears that the endoderm identities emerging and self-organising within Gastruloids to form the core FOXA2+ epithelial domain observed by immunostaining mature endoderm identities corresponding to the entire length of the embryonic gut tube, albeit with low representation of midgut (stomach, pancreas, liver) cells. Notably, Gastruloid-endoderm matures a strong anterior foregut character (corresponding to the pharyngeal pouch endoderm *in vivo*), a finding most recently echoed in gastruloids obtained from zebrafish cells (Cheng et al. (2021), where the only endoderm found was pharyngeal endoderm).

Finally, we verified whether the expression of the markers identified in the single-cell dataset was correctly patterned along the AP axis of the Gastruloid. That is, whether the variety of cell identities uncovered in the single cell dataset are intermingled throughout the core of the Gastruloid, or they are rather spatially segregated at the correct position along the AP axis of the Gastruloid as is the case for their *in vivo* embryonic counterparts along the gut tube. Immunostainings for markers such as TBra and Cdx2 indeed shows positive cells to be restricted to the posterior end of the endoderm primordium (refer back to Figure 4C and Figure 5D). To test whether markers of anterior endoderm populations similarly localise to the anterior of the primordium, we performed Hybridisation Chain Reaction (HCR) against the foregut marker Pax9 (Figure 8D). Indeed, *Pax9* is known to be expressed within the pharyngeal foregut endoderm (Peters et al., 1998) and is highly expressed in the Gastruloid-”mature endoderm” cluster (Figure 8D). For this foregut marker too, expression is consistently restricted to the anteriormost extremity of the endoderm primordium. It thus appears that both posterior and anterior markers can be recovered at their expected position within the endoderm primordium of late Gastruloids.

## Discussion

Cells expressing endodermal markers, and gene expression patterns consistent with endodermal identities, have been described in several previous and current Gastruloid studies (Anlaş et al., 2021; Beccari et al., 2018; Olmsted & Paluh, 2021; Pour et al., 2019; Turner et al., 2017; van den Brink et al., 2020, 2014; Veenvliet et al., 2020). These descriptions have however often not taken centre stage (with the exception of (Pour et al., 2019), see later). Of notice is the fact that FoxA2 and Sox17, indeed classical endoderm markers, are also expressed in other embryonic cell types at the same developmental stages where the endoderm and the gut tube are specified. As such, while detecting either of these markers in the embryo may exclude non-endodermal identities given the spatial and temporal context of the observation, the same cannot be said in Gastruloids, where the full extent of the cell types generated is still under characterisation (van den Brink et al., 2020), and where temporal alignment with *in vivo* developmental stages is uncertain (Beccari et al., 2018). Accordingly, FOXA2 marks the endoderm just as much as the neural floorplate and the notochord; TBra marks posterior hindgut as well as posterior epiblast, notochord, and neuromesodermal progenitors; and SOX17 marks both endoderm and endothelial progenitors (Choi et al., 2012). Regardless, original descriptions of Gastruloids were indeed already describing polarised emergence of endodermal cells expressing both FOXA2 and SOX17 (van den Brink et al., 2014). A compact FOXA2+ domain was thus seen to cluster at the posterior of late stage Gastruloids, with SOX17+/TBra-cells occupying this very FOXA2+ domain and internalising within epithelial vesicles. We presently cannot reconcile this posteriormost pattern of expression with what we describe here, but we do notice that in those cell lines where the CDH1 primordium does not extend throughout the length of the aggregate it segregates as a compact mass at the posterior (data not shown, but see FGF4-treated deficient Gastruloids in Gharibi et al. (2020)). The described invagination of CDH1+/SOX17+ cells may however indeed explain the surface continuity of the posterior CDH1 primordium that we see in the neck region of some of our Gastruloids.

Even more complete Sox17 patterns have been further described in (Turner et al., 2017), where the use of a reporter highlights the formation of *Sox17+* midline, tubular-shaped patterns in elongating gastruloids. The study likens these cells to ventral endodermal cells of the E8.5 mouse embryo. Based on our result, and the extensive SOX17 positivity of the CDH1 primordium we describe here, we would expect the reporter line used in (Turner et al., 2017) to equally give rise to an internal endodermal primordium. Interestingly what we here infer by immunostaining seem to be consistent with the early *Sox17* dynamics described in the paper: Sox17+ cells emerging towards the anterior pole of the early Gastruloid and then expanding to occupy a relatively more posterior domain.

SOX17+ cells were also identified in (Beccari et al., 2018), and described as forming tubular structures based on DAPI counterstaining. More importantly, the publication describes gene expression dynamics associated with advanced endoderm maturation: the early upregulation of markers such as Gsc and Cdx2, upregulation of Cer1 during elongation (and which indeed appear in the “early endoderm” cluster discussed here), and expression of gut endoderm markers during later development (Nedd9, Sorcs2, Pax9, Pyy, Shh, Krt18). *In situ* hybridisation patterns for some of these markers are again consistent with the presence of an internal endodermal structure. Still, detailed spatial characterisation is lacking, and the Gastruloid remains framed as a mainly mesenchymal neuromesodermal aggregate. Validation of the maturation of endodermal identities can also be found in the single-cell transcriptomics data generated by (van den Brink et al., 2020), which detect a cluster of cells postulated to represent definitive endoderm as expressing markers Epcam, Col4a1, Sox17. All of these markers are indeed recovered within the endoderm clusters considered here.

Most recently, tubular structures populated by FOXA2+ and some SOX17+ and TBra+ cells have also been described in (Veenvliet et al., 2020). Associated with such endoderm-like compartment, the authors also observe Primoridal-Germ-Cells (DPPA3+ cells) which indeed migrate along the gut tube during *in vivo* development. Our results would seem to suggest that embedding of the Gastruloid in extracellular matrix components (Veenvliet et al., 2020) is however not necessary to observe endoderm morphogenesis, and that this tissue may actually already organise its own extracellular matrix environment. Unlike mesoderm thus, which seems to require *in vitro* ECM cues for productive mesenchymal to epithelial transition (van den Brink et al., 2020; Veenvliet et al., 2020), morphogenesis of the endoderm appears to be intrinsic to the tissue. Accordingly, Olmsted & Paluh (2021) recently reported the formation of an analogous epithelial central tube in human gastruloids under shaking culture (i.e. without embedding). This so-called “primitive gut tube” expresses classic endodermal markers (*FoxA2, Sox17, Gata6, Sox2, Cdh1*, and *Epcam*), as well as markers of more specialised cell types such as *Lgr5, Nkx2*.*1, Lyz*, and *Vil1* over 16 days of culture. It is not stated whether these markers are patterned along the tube.

The earliest steps of endoderm emergence in Gastruloids have been attentively detailed by (Pour et al., 2019), who elegantly used a TBra/Sox17 double reporter cell line to document expression of these two markers in a temporal sequence consistent with what we describe in this study. *Sox17*+ cells emerge around 1 day after CHIR exposure, intermingled within a field of TBra+ cells. While the authors also see strong association between *Sox17* expression and epitheliality (CDH1 positivity), they do seem to identify a stage where such Sox17+ cells are CDH1-, and indeed favour an interpretation that supports a mesendodermal origin of endodermal precursors. In our observations, Gastruloids start in a epithelioid configuration where cells have mesenchymal character but express CDH1, and in which TBra+, FOXA2+, and SOX17+ cells thus all emerge in a CDH1+ context. Early endoderm dynamics have since also been reported in Anlaş et al. (2021), which used HCR to show early anterior segregation og *Foxa2*+ cells in steps and relative timings consistent with what shown here by immunostaining and live imaging.

The observations we report here bring centre stage the question of the epithelial character of endodermal precursors, and its link to the fate (both in terms of identity and of location) of these cells (Ferrer-Vaquer et al., 2010; Nowotschin et al., 2019a; Viotti et al., 2014). We expand on previous Gastruloid studies by taking into consideration multiple markers of endodermal identity and documenting their dynamics in relation to one another, and notably by also tracking these cells over time as they undergo morphogenesis. We further also describe sorting of these cells into a complex primordium with maturing epithelial architecture and coarsely patterned domains of gene expression along the AP axis of the organoid. By transcriptional comparison with relevant embryonic datasets (Nowotschin et al., 2019b), we highlight the emergence of cell identities corresponding to the entire length of the gut tube, with notable representation of anterior foregut, midgut, and posterior hindgut types, and show these to be patterned along the AP axis of the system.

At this point, our observations are consistent with a model whereby SOX17+ cells never leave their epithelial environment (initially, the “epiblast”-like domain) and do not need to transition through a mesenchymal state, at least not through classic Snai1-mediated EMT. We speculate that, if SOX17+ endodermal cells are not being directly specified within the FOXA2+ epiblast, the SOX17+ cells that we incongruously see in this compartment are a result of these cells sticking to neighbours with the same epithelioid character. While in the embryo the isotropic relocalisation of CDH1 associated with egressed endodermal cells might be compatible with segregation from both epiblast (columnar epithelium) and visceral endoderm (squamous epithelium), and indeed reintegration of Sox17+ cells requires re-polarisation, the situation is different in Gastruloids. In our system, FOXA2+ and SOX17+ cells are emerging not in a polarised columnar epithelium, but in a context that already displays isotropic CDH1 localisation, such that these cells may remain stuck in their original compartment just by virtue of homotypic interactions. Interestingly, and during preparation of this manuscript, the use of a Cdh1 live reporter by (Hashmi et al., 2020) has shown fragmentation and early sorting dynamics consistent with what we show here by immunostaining, and the authors explain these sorting dynamics by differences in interfacial tension across cell types. Later in development we here further observe an expansion of the endodermal population and its internalisation within the core of the Gastruloid as the surface of the aggregate start being populated by an increasing number of mesodermal cells. Interestingly, the relative position of different epithelial populations may here again be explained by the expression of different combinations of cell-surface adhesion molecules, a common sorting mechanisms (Toda et al., 2018) that sees here some support from the biased EpCAM distribution within our CDH1 primordium, enriched in the domains occupied by endodermal cells.

Our identification of a maturing epithelial structure throughout late Gastruloid development, contrasting with the overlaying mesenchymal mesodermal tissues enveloping it, reframes expectations regarding the extent to which fate and morphogenesis can spontaneously arise *in vitro*. While gastruloids have traditionally been pictured as aggregates of fates without corresponding organisation, we start to see increasing examples where such missing morphogenesis does occur. As already shown by (Bérenger-Currias et al., 2020; van den Brink et al., 2020; Veenvliet et al., 2020) this can be transformatively brought about by the addition of diluted ECM components or extraembryonic cell types. We here show that differences in epithelial identities between emerging cell types may already be sufficient to generate simple architectures, and that complex epithelia may spontaneously organise *de novo* in Gastruloids. Still, the extent to which the elaborate morphogenesis seen here is cell-line specific remains to be defined. Current literature contains examples of Gastruloids from a variety of cell lines and mouse backgrounds that do indeed originate such internal epithelial primordia (Beccari et al., 2018; Rossi et al., 2019) (or where we infer such primordium to be what described (Gharibi et al., 2020; Turner et al., 2017; Veenvliet et al., 2020)), as well as of Gastruloids where such structures do not seem to appear (van den Brink et al., 2020, 2014) and where CDH1+ endodermal tissue segregates to the posteriormost tip of the aggregates instead. We favour the hypothesis that these differences likely correlate to intrinsic cell-line biases in core signalling pathways, and/or on the very degree of epitheliality maintained by these cells by the time they are exposed to CHIR (in turn possibly relating to differences in the 2D culture conditions of these cells). A systematic assessment of the cell-line specific differences in these key parameters remains to be performed, yet this hypothesis seems to be supported by the fact that pretreatment with different signalling factors can allow/prevent formation of the primordium as shown in Gharibi et al. (2020). As is the case with the cell line used in this study (Figure 5), we nonetheless expect all Gastruloids generated from a given cell line to consistently produce the same endoderm phenotype in all Gastruloids generated. Regardless, the stratified nature of endodermal and mesodermal tissues we here observe is at least broadly comparable to the configuration of endoderm and mesoderm in the embryo, and it is interesting to speculate that this interfacing may favour the development of more advanced cell fates by reciprocal signalling interactions, as*in vivo* (Bardot & Hadjantonakis, 2020; Han et al., 2019).

As per the classical paradigm offered by *in vitro* developmental models (Shahbazi & Zernicka-Goetz, 2018), we believe that the extent to which Gastruloid endoderm development matches *in vivo* gut tube development will highlight important developmental principles in both systems. For now, our observations at early stages of Gastruloids development support the existence of endodermal progenitors that do not transition through a TBra+ state and that do not necessarily undergo classic EMT, something debated in the field and for which there is not yet conclusive evidence for in the mouse embryo, but consistent with what seen in other embryonic models (Nowotschin et al., 2019a). Regarding the abnormal epiblast-retention of endodermal cells in Gastruloids, which we explain by the incongruous absence of a starting epithelial architecture in the system, we wonder whether a similar conjuncture would be seen in mutant embryos in which the epiblast does not maintain apico-basal polarity and where epiblast CDH1 may thus be also already isotropically distributed. FOXA2+ and SOX17+ cells that populate the Gastruloids at 120h further pattern according to the anterior-posterior cues of the aggregate, which thus produces anterior foregut (pharyngeal) identities at the anterior, and hindgut identities at the posterior. We thus postulate that Gastruloids could be a valid source for the isolation and further differentiation of specific endodermal identites which may be otherwise more difficult to differentiate *in vitro* (e.g. thymus from anterior foregut tissue).

## Materials and methods

### Cell culture

mESCs (“SBR” Sox1/TBra double reporter cell line described in (Deluz et al., 2016); CRG8 cells of 129P2 background (RRID:CVCL_3987, Mountford et al. (1994)); or “TFoxA2” FoxA2/TBra double reporter cell line described in (Yang, 2015); E14 cells of 129P2 background (RRID:CVCL_C320, Fehling et al. (2003); Hooper et al. (1987))) were cultured in tissue-culture-treated, non-gelatinised, 6well plates, in 10%Serum Medium with added 2i and LIF. Cells were split every third day, by washing in PBS-/-, adding Accutase for ∼3min RT, and collecting the resulting cell suspension in a clean centrifuge tube. The Accutase of the cell suspension was then diluted out 1:10 in 10% Serum Medium, and cells were pelleted by centrifuging 200xg, 4min, 4°C. After aspirating out the supernatant, the pellet was resuspended in 1mL 10% Serum Medium to a single cell suspension, and cell density was counted with a haemocytometer. Around 65000-75000 cells were transferred to a new well with 2mL pre-equilibrated 10%Serum Medium (6750-7800 cells/cm2). Cells were then left in a humidified incubator, 37°C, 5% CO_2_, until use or until further splitting 3 days later. NOTE: In most cases, splitting was coupled to Gastruloid generation. In those cases, the cell pellet was resuspended in N2B27 rather than 10% Serum Medium + 2i and LIF. A complete, step-by-step, protocol is available at:https://dx.doi.org/10.17504/protocols.io.7xbhpinRecipes: 10% Serum Medium: 86.8% DMEM, high glucose, with GlutaMAX (L-Alanyl-Glutamine, final concentration: 3.45mM), 10% ES-grade Foetal Bovine Serum, 100U/mL Penicillin, 100ug/mL Streptomycin, 0.1mM Non Essential Amino Acids, 1mM Sodium Pyruvate, 0.1mM beta-mercaptoethanol 10% Serum Medium + 2i and LIF: add 3uM CHIR99021, 1uM PD0325901,and 100u/mL LIF.

### Gastruloid generation

mESCs were washed in PBS-/-, detached from adherent culture with Accutase (∼3min, RT), and collected in a centrifuge tube. The Accutase in the cell suspension was then diluted out 1:10 in 10% Serum Medium, and cells were pelleted by centrifuging 200xg, 4min, 4C. The supernatant was removed, and the pellet was washed by resuspension in 10mL PBS-/-. Cells were re-pelleted by centrifuging 200xg, 4min, 4C and washed once more in 10mL fresh PBS-/-. After re-pelleting once more (200xg, 4min, 4C), the pellet was dissociated as a single-cell suspension in 1mL N2B27 Medium. Cells were counted with a haemocytometer, and, for each plate of Gastruloids made, 37500 cells (SBR line) or 93750 cells (TFoxA2 line) were transferred to 5mL fresh N2B27 (7.5 cells/uL or 18.75 cells/ul final concentration, respectively). The cell suspension was distributed as 40uL droplets (=300 SBR cells/droplet, or 750 TFoxA2 cells/droplet) in wells of a U-bottomed, low-adhesion, 96 well plate, and the plates were left for 120h in a humidified incubator, 5% CO_2_, 37°C. At 48h after plating, 150uL of 3uM CHIR99021 N2B27 were added to each well, and this solution was substituted with fresh N2B27 (no CHIR) every 24h after that. A step-by-step detailed protocol is available at: https://dx.doi.org/10.17504/protocols.io.9j5h4q6 Recipes: 10% Serum Medium: 86.8% DMEM, high glucose, with 3.97mM GlutaMAX (L-Alanyl-Glutamine, final concentration: 3.45mM), 10% ES-grade FBS, 100U/mL Penicillin, 100ug/mL Streptomycin, 0.1mM Non Essential Amino Acids, 1mM Sodium Pyruvate, 1mM beta-mercaptoethanol. N2B27: 47.4% Neurobasal Medium, 47.4% DMEM/F-12, with 2.50mM GlutaMAX (L-Alanyl-Glutamine, final concentration: 1.18mM), 1mM GlutaMAX Supplement (total concentration: 2.18mM), 100U/mL Penicillin, 100ug/mL Streptomycin, 0.1mM Non Essential AminoAcids, 1mM Sodium Pyruvate, 1mM beta-mercaptoethanol, 1% B27Supplement, serum-free, 0.5% N-2 Supplement.

### Gastruloid immunostaining

Gastruloids were collected at every given timepoint, washed in PBS-/-, and fixed in 4% PFA in PBS-/-, for 2h, 4C, on a low-speed or-bital shaker; or 45min, RT, static. Gastruloids were then washed in PBS+FT (PBS-/-, 10% ES-grade Foetal Bovine Serum, 0.2% Triton-X100), and blocked and permeabilisedin PBS+FT for 1h, RT, static. Primary antibody solutions were then prepared in PBS+FT, with 2ug/mL DAPI. Samples were stained overnight, 4C, on a low-speed orbital shaker. Similarly, secondary antibody solutions were prepared in PBS+FT, 2ug/mL DAPI, and samples were stained overnight, 4C, on a low-speed orbital shaker. Gastruloids were mounted in Fluoromount-G mounting medium (no spacers), and slides kept at 4C long term. All antibody solutions were washed away after incubation by washes in PBS+FT. A detailed, step-by-step protocol is available at: https://dx.doi.org/10.17504/protocols.io.7tzhnp6. Secondary antibodies used were all from Thermo Fisher Scientific: donkey anti-mouse IgG Alexa 647 (CAT#A-31571, RRID:AB_162542); donkey anti-rabbit IgG Alexa 488 (CAT#A-21206, RRID:AB_2535792), Alexa 568 (CAT# A-10042, RRID:AB_2534017), or Alexa 647 (CAT# A-31573, RRID:AB_2536183); donkey anti-rat IgG Alexa 488 (CAT#A-21208, RRID:AB_2535794), goat anti-rat Alexa 568 (CAT#A-11077, RRID:AB_2534121), or Alexa 647 (CAT#A-21247, RRID:AB_141778); donkey anti-goat IgG Alexa 488 (CAT#A-11055, RRID:AB_2534102), or Alexa 568 (CAT#RRID:AB_2534104). Details about the primary antibodies used are provided as a supplementary .csv file.

### Gastruloid FACS

TFoxA2 Gastruloids were grown according to the protocol described above, and two/four 96well plates of Gastruloids at each timepoint were used and processed for FACS. Briefly, Gastruloids were collected at every given timepoint, washed in PBS-/-, and digested 8min, 37C, in Digestion Solution (Collagenase IV [3mg/mL],Dispase [4mg/mL], DNAseI [100ug/mL], in PBS). Working on ice, the cell suspension was then strained through the filter cap of a FACS tube, and an excess of cold Staining Buffer (10%ES-FBS, Pen-Strep [100U/mL], EDTA [1mM], in PBS) was added to stop the digestion. Cells were then stained with DAPI ([0.2ug/mL] DAPI, in Staining Buffer), 10min, 4C, fixed in 2%PFA, 4C, 10min, and stored in Staining Buffer, 4C, in the dark, until use. Standard 2D cultures of TFoxA2 mESCs and RFP+ mESCs were used as negative and positive references, respectively. These were detached in Accutase, 4min, RT, and DAPI-stained and fixed as done for filtered Gastruloid cells and as described above. GFP BrightComp eBeads™ (Invitrogen/Thermo Fisher Scientific, CAT#A10514) were used as GFP+ positive reference, according to manufacturer protocol (1 drop of beads resuspended in 1mL Staining Buffer). A step-by-step detailed protocol is available at: https://dx.doi.org/10.17504/protocols.io.bvgrn3v6 Samples were analysed on a Becton Dickinson LSR-Fortessa™ Flow Cytometer, with optical configuration 355nm[450/50], 488nm[530/30], 561nm[585/15], using BD FACSDivaTM software, with applied compensation. Exported FCS files were analysed in RStudio (ggcyto library, Van et al. (2018), and flowCore library, Ellis et al. (2021)). The annotated notebook, with a step by step walkthrough the entire analysis pipeline, is available at: https://doi.org/10.5281/zenodo.4894122

### Gastruloid Hybridisation Chain Reaction (HCR)

Gastruloids were collected at every given timepoint, washed in PBS-/-, and fixed in 4% PFA in PBS-/-, overnight, 4C, on a low-speed orbital shaker. Gastruloids were then washed in PBS-/-, and then dehydrated in a graded series of methanol-PBST solutions (0%-100%, 25%-75%, 50%-50%, 75%-25%, 100% methanol). Gastruloids were then stored in 100% methanol, −20C, until use (and at least overnight). When needed, Gastruloids were rehydrated in a graded series of methanol-PBST solutions (100%-0%, 75%-25%, 50%-50%, 25%-75%, 100% PBST), digested in 25ug/mL Proteinase K in PBST, 4min, RT, washed in PBST, and re-fixed in 4%PFA in PBS-/-for 20min, RT. For the probe hybridisation step, samples were washed in PBST, pre-incubated 1h30min in warm Probe Hybridisation Buffer, 37C, and then incubated for 16-20h with 4pmol of odd HCR probes and 4pmol of even HCR probes mixed in Probe Hybridisation Buffer, 37C. For the amplification step, samples were washed in warm Probe Wash Buffer, 37C, further washed in RT 5XSSCT, and then left to incubate for 16-20h with 48pmol of hairpin 1 and 48pmol of hairpin h2 (for each colour used) mixed in Probe Amplification Buffer with 2ug/mL DAPI, RT. Each hairpin was heated to 95C for 1min30s, and snap-cooled at RT for at least 30min before use. After amplification, samples were incubated for 1h15min in 5XSCCT with 2ug/mL DAPI, washed in 5XSCCT, and then mounted on microscope coverslips in Fluoromount G mounting medium. A step-by-step detailed protocol is available at: https://dx.doi.org/10.17504/protocols.io.bcwfixbn The sequences of the *Pax9* probe set used (coupled to amplifier B5), is provided as a supplementary file.

### Processing of scRNAseq datasets

All data analysis was done on R, with the Seurat v4.0 library (Hao et al., 2020) **Gastruloid dataset:** scRNAseq data corresponding to Gastruloids spanning timepoints t=96-168h was taken from Rossi et al. (2019). Raw count matrices for both batches of each timepoint and for different timepoints were merged, and filtered based on the following quality control parameters: *number of Unique molecular identifiers* > 10000, *number of Genes* > 2000, *Complexity* > 0.75, *Percentage of mitochondrial genes* < 15%. Genes expressed in 0 or less than 5 cells of the dataset were discarded. The data underwent normalisation, variance stabilisation, and differences due to mitochondrial content and cell cycle phase were regressed out via Seurat’s *NormalizeData* and *SCTransform* functions. Data from different timepoints was integrated using *SelectIntegrationFeatures* on the top 3000 genes, *FindIntegrationAnchors*, and *IntegrateData*. PCA and UMAP (on the first 40 dimensions) were then calculated through *RunPCA* and *RunUMAP*, respectively. Clustering was done via the functions *FindNeighbours* and *FindClusters*, by a shared nearest neighbor (SNN) modularity optimization based clustering algorithm on a SNN graph based on the 20 nearest neighbours (default). A resolution of 0.8 was chosen to proceed with analysis. Cluster identities were assigned based on the patterns of expression of selected marker genes, along with an analysis of the genes marking each cluster (*FindaAllMarkers* function, limiting testing to genes which showed, on average, at least 0.25-fold difference (log-scale) between the two groups of cells; default). Top genes were ranked based on the difference between their pct.1 and pct.2 values (i.e. between the percentage of cells expressing a given gene in the cluster of interest versus in the other clusters combined). To find markers differentiating close clusters, the function *FindMarkers* was used instead. The annotated notebook, with a step by step walkthrough the entire analysis pipeline, is available at: https://github.com/StefanoVianello/Endoderm_scRNAseq, “Gastruloid_scRNAseq_preprocessing_RNotebook.Rmd”.

### Gut endoderm dataset

scRNAseq data corresponding to Gut endoderm cells at Embryonic Day 8.75 was taken from Nowotschin et al. (2019b). The raw count matrix was imported as a Seurat object and filtered based on the following quality control parameters: *number of Unique molecular identifiers* > 5000, *number of Genes* > 3000, *Complexity* > 0.75, *Percentage of mitochondrial genes* < 20%. Genes expressed in 0 or less than 5 cells of the dataset were discarded. The data underwent normalisation, variance stabilisation, and differences due to mitochondrial content and cell cycle phase were regressed out via Seurat’s *NormalizeData* and *SCTransform* functions. PCA and UMAP (on the first 30 dimensions) were then calculated through *RunPCA* and *RunUMAP*, respectively. *In vivo* and *in vitro* data was integrated using *SelectIntegrationFeatures* on the top 3000 genes, *FindIntegrationAnchors*, and *IntegrateData*. PCA and UMAP (on the first 40 dimensions) were then calculated through *RunPCA* and *RunUMAP*, respectively. Clustering was done via the functions *FindNeighbours* and *FindClusters*, by a shared nearest neighbor (SNN) modularity optimization based clustering algorithm on a SNN graph based on the 20 nearest neighbours (default). A resolution of 0.8 was chosen to proceed with analysis. Cluster identities were assigned based on the patterns of expression of selected marker genes, along with an analysis of the genes marking each cluster (*FindMarkers* function, limiting testing to genes which showed, on average, at least 0.25-fold difference (log-scale) between the two groups of cells; default). Top genes were ranked based on the difference between their pct.1 and pct.2 values (see explanation above). The annotated notebook, with a step by step walkthrough the entire analysis pipeline, is available at: https://github.com/StefanoVianello/Endoderm_scRNAseq, “GutTube_scRNAseq_preprocessing_RNotebook.Rmd”.

### Alignment of Gastruloid and Gut tube cells

Cells corresponding to endoderm clusters were subsetted from the Rossi et al. (2019) dataset, processed as described above. The identification of the endoderm clusters is justified in this manuscript. Cells corresponding to the embryonic gut tube were subsetted from the Nowotschin et al. (2019b) dataset, processed as described above. The gut tube cluster is the biggest cluster in the dataset, and its endodermal identitiy was confirmed based on the expression of classic endodermal markers. The count matrices of the two subsetted datasets (*in vitro* endoderm and *in vivo* endoderm) were merged into a single object and processed according to standard pipeline. Genes expressed in 0 or less than 5 cells of the dataset were discarded. The data underwent normalisation, variance stabilisation, and differences due to mitochondrial content and cell cycle phase were regressed out via Seurat’s *NormalizeData* and *SCTransform* functions. PCA and UMAP (on the first 30 dimensions) were then calculated through *RunPCA* and *RunUMAP*, respectively. Clustering was done via the functions *FindNeighbours* and *FindClusters*, by a shared nearest neighbor (SNN) modularity optimization based clustering algorithm on a SNN graph based on the 20 nearest neighbours (default). A resolution of 0.8 was chosen to proceed with analysis. Cluster identities were assigned based on the patterns of expression of selected marker genes, along with an analysis of the genes marking each cluster (*FindaAll-Markers* function, limiting testing to genes which showed, on average, at least 0.25-fold difference (log-scale) between the two groups of cells; default). Top genes were ranked based on the difference between their pct.1 and pct.2 values (see explanation above). To find markers differentiating close clusters, the function *FindMarkers* was used instead. The annotated notebook, with a step by step walkthrough the entire analysis pipeline, is available at: https://github.com/StefanoVianello/Endoderm_scRNAseq, “Endoderm_comparison_RNotebook.Rmd”.

### Gastruloid imaging and image processing

Bright-field images of Gastruloids were taken on either a Nikon Ti inverted spinning-disk microscope (for the series in Figure 2A), or an Olympus CellR inverted widefield microscope (for the image on Figure 4B, UPLAN S APO 10x/0.40 air objective, CCD Grayscale Hamamatsu ORCA ER B7W Camera; Olympus XCellence software for data capture). Both microscope setups had CO2 and temperature control (37°C and 5% CO_2_). Live imaging of TFoxA2 reporter Gastruloids was done on the Olympus CellR inverted wide-field microscope described above, with acquisition every 30min (5 z-slices, 27.5um spacing) Immunostained Gastruloids were imaged on a Zeiss LSM700 inverted confocal microscope (Plan-Apochromat 20x/0.80 air objective, motorized stage, LED Lumencor SOLA Illumination, CCD Grayscale Axiocam MRm (B/W) Camera; ZEN 2009 software for data capture) or on a Zeiss LSM780 inverted confocal microscope (for the CDH1 projection in Figure 4A, Plan-Apochromat 20x/0.80 air objective, motorized stage). Images were opened, stitched, and processed for publication (LUT assignment, channel display, min and max intensity thresholding based on no-primary control) using the Fiji ImageJ distribution (Rueden et al., 2017; Schindelin et al., 2012), and the “Grid/Collection Stitching” plugin therein (Preibisch et al., 2009). The depth-coded reconstruction in Figure 4A was generated using the “Temporal-Color Code” (https://imagej.net/Temporal-Color_Code) function. The blue, orange, and purple LUTs used throughout the figures were designed by Christophe Leterrier (https://github.com/cleterrier/ChrisLUTs, “BOP” palette).

### Quantification of AP patterning

Batch quantification of immunostaining signal intensity along the AP axis of the Gastruloids was performed through a custom processing pipeline available as a Jupyter notebook at https://doi.org/10.5281/zenodo.4899121 and outlined as follows (step-by-step walk through provided in the notebook itself). The pipeline takes two inputs: i) the multichannel raw image resulting from the scan of an entire microscope slide of immunostained and mounted Gastruloids (here acquired on a GE Healthcare IN Cell Analyzer 2200 automated microscope) and ii) hand-traced line coordinates defining the central axis of each Gastruloid on the slide (starting from the posterior). At early timepoints where the posterior of the Gastruloid is not distinguishable morphologically, the area of TBra polarisation is to be used instead. The script then subdivides each interval of the line ROI provided into *n* finer intervals of equal length (thus avoiding to have to manually draw a line with high number of points; here *n*=10), and for each point along the line it defines a non-overlapping polygon mask covering an area of thickness *N* (here *N*=500px) across the line and whose lateral edges are orthogonal to the line itself at each side of the point. Having computed the mask, the script then assigns the total signal intensity recovered in the area to the point of the line ROI around which the polygon was constructed, thus effectively assigning signal intensities to points that can be ordered along an x-axis. These raw values are then normalised by the number of cells in the area (using the DAPI nuclear intensity as a proxy) and both position along the length of the Gastruloid and signal intensity are normalised to the absolute length of the Gastruloid and to the maximal DAPI-normalised intensity value. The script outputs lineplots and scatterplots for each Gastruloid analysed, summary lineplots and scatterplots with collated data of all gastruloids analysed, and the tabulated raw data for re-use.

## Supporting information

Supplementary Table (Primary Antibodies)

Supplementary Table (markers of Gastruloid clusters)

Sequence of Pax9 HCR probes

Video 1 (FoxA2/TBra Gastruloid timelapse)

## ACKNOWLEDGEMENTS

We thank the Lickert lab (Helmholtz Zentrum München) for sharing the FoxA2/TBra double reporter cell line used. We would like to acknowledge the work and support provided by the staff of the following EPFL core facilities: the BioImaging and Optics Platform (BIOP), the Biomolecular Screening Facility (BSF), the Histology Core Facility, the Flow Citometry Core Facility (FCCF). We also want to acknowledge the work of the entire staff of the Glassware Washing facility (Laverie), as well as EPFL cleaning staff. This work would have not been possible without their contribution. The pipeline designed to quantify AP patterning in Gastruloids was constructed in collaboration with Paul Gerald Layague Sanchez and Arianne Bercowsky-Rama, whom we sincerely thank. We would also like to acknowledge our lab manager Stephanie Boy-Rottinger for her heartfelt support throughout. In addition, we are thankful to Ricardo Henriques for kindly sharing the template that was used to format this preprint. We also thank Alfonso Martinez Arias and Heiko Lickert for valuable feedback on the first uploaded bioRxiv version of this manuscript. This research was funded by a Sinergia grant (CRSII5_189956) from the Swiss National Science Foundation and Ecole Polytechnique Fédérale de Lausanne (EPFL).

---

**Fig. S1.**
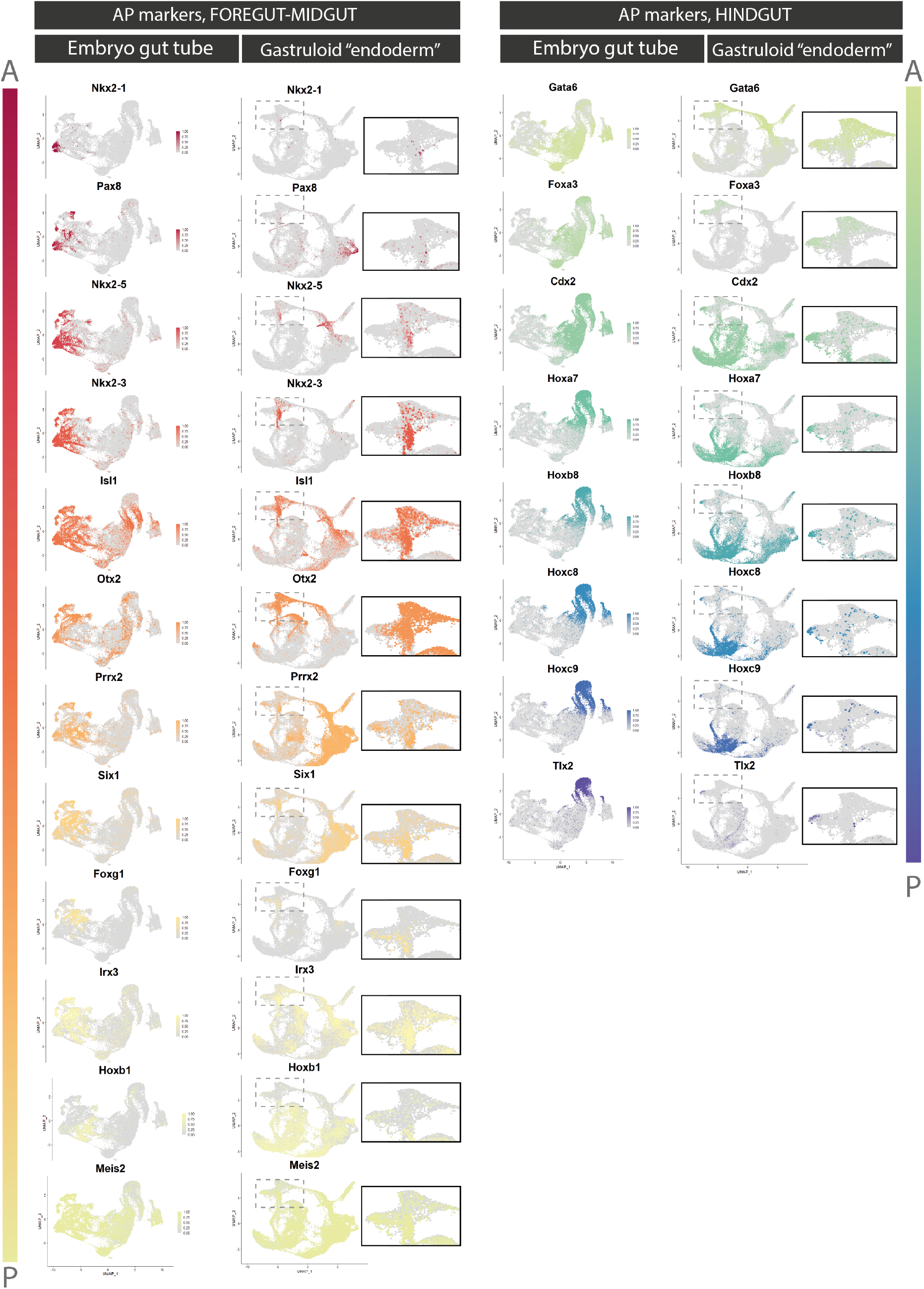
Expression pattern of gut endoderm Anterior-Posterior markers in *in vivo* and *in vitro* datasets. Markers of AP position along the embryonic gut tube (Nowotschin et al., 2019b) are plotted from anteriormost foregut (top, *Nkx2-1*), to posteriormost hindgut (bottom, *Tlx2*). For each gene, the expression pattern is shown for both the embryonic dataset (E8.75 gut tube, Nowotschin et al. (2019b); **left**) and the Gastruloid dataset (SBR Gastruloids, Rossi et al. (2019), **right**). Validating the reprocessing of the embryonic dataset, increasingly posterior markers define continuous domains from one extremity of the UMAP to the other. For the Gastruloid dataset, a inset focusing on the two “endoderm” clusters is also provided. A = Anterior, P = Posterior.

